# Reelin Deficiency Alters the Directional Orientation, Apical Localization and Length of Primary Cilia in Principal Neurons in the Cerebral Cortex

**DOI:** 10.1101/2025.08.24.672028

**Authors:** Sumaya Akter, Soheila Mirhosseiniardakani, Yuto Takekoshi, Liyan Qiu, Kevin Baker, Kevin Jiang, Mark Lyon, Mitsuharu Hattori, Xuanmao Chen

## Abstract

The primary cilia of pyramidal neurons in inside-out laminated regions are oriented predominantly toward the pia, a pattern reflecting reverse soma movement during postnatal neurodevelopment. However, the mechanisms that regulate the directional cilia orientation remain unknown. Here we show that the primary cilia of neocortical pyramidal neurons are localized at the base of the apical dendrites and aligned distally and opposite to the axon initial segment (AIS). However, this pattern is not observed in atypical pyramidal neurons in the deep neocortex, excitatory neurons in non-laminated regions, or interneurons, where cilia are randomly positioned around the nuclei and lack preferred orientation. In Reelin-deficient mice (*reeler*), the directional orientation and apical location of cilia in late-born neocortical and CA1 neurons are altered. However, the initial impairments are partially corrected during early postnatal development. In contrast, Reelin deficiency disrupts the directional orientation and apical localization of primary cilia in early-born neocortical neurons and principal neurons in evolutionarily conserved laminated regions, where lack postnatal correction. Reelin deficiency also increases the cilia length in principal neurons across the cerebral cortex at a developmental stage when cilia length stabilizes in wild-type mice. In contrast, cilia elongation is not observed in interneurons, GFAP-positive astrocytes, or excitatory neurons in non-laminated *reeler* regions. Together, Reelin regulates the directional orientation, apical localization, and length of primary cilia in principal neurons in the cerebral cortex. This study underscores the cilium as a key apical domain, particularly prominent in late-born pyramidal neurons.

## Introduction

The extracellular matrix protein Reelin is a crucial determinator directing embryonic neuronal migration of principal neurons and lamina formation in the cerebral cortex (Alexander et al., 2023; Hattori, 2025; Jossin, 2020). Reelin exerts its effects through binding to two lipoprotein receptors, ApoER2 and VLDLR, initiating a signaling cascade involving the phosphorylation of the adaptor protein Dab1 (Frotscher et al., 2009; Trommsdorff et al., 1999). By modulating cytoskeleton dynamics, stabilizing leading process, and regulating soma translocation (Jossin, 2020), Reelin regulates the establishment and maintenance of apical-basal polarity of principal neurons in the cerebral cortex (Kupferman et al., 2014; Matsuki et al., 2010). Disruption of Reelin signaling, as seen in Reelin null mutant *reeler* (D’Arcangelo et al., 1995), results in abnormal neuronal morphology, apical-basal misorientation, and profoundly disorganized cortical formations. These changes are associated with numerous downstream consequences, including synaptic dysfunction and severe neurological disorders such as lissencephaly, ataxia, and schizophrenia and neurodegeneration (Jossin, 2020). While Reelin is known to guide embryonic neuronal migration and cortical layering (Alexander et al., 2023; Jossin, 2020; Katsuyama and Hattori, 2024), its impacts on postnatal neuronal positioning and subcellular neuronal polarity are not well understood.

Primary cilia are centriole-derived organelles that are expressed in most mammalian cells (Wilsch-Bräuninger and Huttner, 2021; Yang et al., 2025). Cilia length varies greatly across brain regions (Wu et al., 2024), developmental stages, day and night rhythm, as well as metabolic or disease conditions (Macarelli et al., 2023; Phua et al., 2019; Sterpka et al., 2020; Tu et al., 2023; Yang et al., 2025). Altered ciliary signaling affects neurodevelopment, leading to various ciliopathies, encompassing cortical malformation, cognitive deficits, and behavioral abnormalities (Green and Mykytyn, 2014; Hasenpusch-Theil and Theil, 2021; Lee and Gleeson, 2011; Singla and Reiter, 2006). During embryonic neurodevelopment, primary cilia coordinate key signaling pathways such as the sonic hedgehog signaling (Bangs and Anderson, 2017; Breunig et al., 2008; Ho and Stearns, 2021) and WNT signaling (Niehrs et al., 2025; Vuong and Mlodzik, 2023), and regulate multiple developmental processes (Guo et al., 2015; Liu et al., 2021; Valente et al., 2014), including cell fate determination (Jang et al., 2016; Yanardag and Pugacheva, 2021) and dorsal-ventral patterning of the neural tube (Pal and Mukhopadhyay, 2015; Sasai and Briscoe, 2012). However, specifically the primary cilia of principal neurons (excitatory, projection, mostly pyramidal neurons) in the cerebral cortex do not emerge during the embryonic stage (Yang et al., 2025). The emanation of primary cilia out of the centrioles, which occurs between P3 to P14 depending on cortical regions, may reflect the dynamic process of principal neuron positioning (Yang et al., 2025). We analyzed cilia length and orientation in the postnatal mouse brain and discovered that after postnatal day 14 (P14), the primary cilia of principal neurons in inside-out laminated cerebral cortical regions, spanning from the hippocampal CA1 region to neocortex, are predominantly oriented toward the pia direction (Yang et al., 2025). The directional orientation of cilia reveals a slow “reverse soma movement” of pyramidal neurons during postnatal positioning (Yang et al., 2025).

Yet, it is unknown what factor controls or regulates the directional cilia orientation in principal neurons in the cerebral cortex. Using Reelin deficient mice (*reeler*) (Curran et al., 1995; D’Arcangelo et al., 1995; Falconer, 1951; Ogawa et al., 1995), we discovered that Reelin deficiency affects directional cilia orientation and cilia localization to the base of apical dendrites of pyramidal neurons. Surprisingly, Reelin deficiency affects early-born neurons and principal neurons in the piriform cortex, hippocampal CA3 and dentate gyrus (DG) more strongly than late-born neurons with respect to cilia orientation, cilia apical localization or apical-basal neuronal orientation. In addition, Reelin deficiency also promotes cilia elongation in principal neurons throughout the cerebral cortex after P14, a developmental stage when primary cilia stabilize in WT mice (Yang et al., 2025). This study highlights primary cilia as apical domains in pyramidal neurons and reveals a previously unrecognized role of Reelin in subcellular neuronal polarity.

## Methods and Materials

### Animals

Mice were housed under a 12-hour light/dark cycle (lights on from 6:00 A.M. to 6:00 P.M.) and provided with standard chow and water ad libitum. Crl:CD-1 mice (RRID: MGI:5652673) were obtained from Japan SLC. *Reeler* mice (B6C3Fe a/a-Reln^rl^ /J, RRID: IMSR_JAX:000235) were obtained from the Jackson Laboratory and backcrossed to the Crl:CD-1 strain. Both male and female mice were used in all experiments. Mice were fixed with 4% paraformaldehyde (PFA) in phosphate-buffered saline (PBS) by intracardiac perfusion. The brains were post-fixed with 4% PFA in PBS overnight before dehydration using 30% sucrose. If not otherwise indicated, experiments were performed using wild-type mice with C57BL/6j genetic background. All animal procedures were approved by the Institutional Animal Care and Use Committee of the University of New Hampshire (Approved Numbers: 191102 and 221202) or by the Animal Care and Use Committee of the Nagoya City University and conducted in accordance with the Institutional Guidelines for Animal Experimentation (Approval Number: 23-001H05).

### Immunofluorescence Staining

Mouse brains were sliced to 30 µm and mounted on gelatin-coated slices for doing immunostaining method. They were washed three times with 0.1 M PBS. Then they were washed in 0.2% Triton X-100 (vol/vol) in 0.1 M PBS three times for 10 minutes at room temperature. Then they were blocked in blocking buffer (0.1M PBS containing 10% normal donkey serum (vol/vol), 2% bovine serum albumin (weight/vol), 0.2% Triton X-100 (vol/vol)) for 1-2 h at room temperature. After that, slices were incubated with primary antibodies overnight at 4 °C in blocking buffer. After washing in 0.1 M PBS three times for 10 minutes, slices were incubated with secondary antibodies for 2 hours at room temperature. Primary antibodies were rabbit anti-AC3 (1:10000 dilution, EnCor Biotechnology Inc., #RPCA AC3) or chicken anti-AC3 (1:1000 EnCor Biotechnology Inc., Cat# CPCA-ACIII), Rabbit anti-Glutaminase I (1:500, Synaptic Systems #456003), rat anti-Ctip2 (1:500, Millipore-Sigma, MABE1045), rabbit anti-Satb2 (1:500, Abcam, ab92446), mouse anti-Ankyrin G (1:500, Synaptic System, #181011), and mouse anti-NeuN (1:500, Millipore-Sigma, MAB377, clone A60). Secondary antibodies were from Donkey conjugated with Alexa Fluor 488, 546, or 647 (Invitrogen, La Jolla, CA). Finally, sections were counter-stained with DAPI (Southern Biotech, #OB010020). A supplemental table lists detailed information on sources of antibodies and their working dilution (Table S1). WT and *reeler* tissue sections used for direct comparisons were processed in parallel using the same antibody solutions and concentrations, incubation times, washing conditions, and secondary antibodies. Corresponding images were acquired using the same confocal settings.

### Confocal Microscopy

Fluorescence images were acquired using a Nikon A1R-HD laser-scanning confocal microscope mounted on a Nikon Ti2 inverted microscope installed in the University Instrument Center. All images were acquired using a Plan Fluor 40× oil-immersion objective (numerical aperture, 1.3; immersion-medium refractive index, 1.515). DAPI and the green, red, and far-red fluorophores were excited using approximately 405-, 488-, 561-, and 640-nm laser lines, respectively. Images were acquired at 1024 × 1024 pixels with a pixel size of 0.310 µm, corresponding to a field of view of approximately 317 × 317 µm. Z-stacks contained 13–24 optical sections acquired at 1-µm intervals, spanning 12–23 µm from the first to the last optical plane. The confocal pinhole was set to 1.2 Airy units. Images were acquired using a unidirectional galvano scan at 1× scanner zoom, scan-speed settings of 0.25 - 0.5, 12-bit depth, and without line averaging. Because different antibody and fluorophore combinations were used across immunostaining experiments, laser power and detector gain and offset were optimized separately for each staining combination but maintained consistently between the experimental groups directly compared within each experiment. Multichannel images were acquired using sequential channel scanning to minimize spectral overlap.

### Cilia Orientation Measurement and Analysis

A custom MATLAB® App, named Cilia Locator (Mirhosseiniardakani et al., 2025), was developed to automatically identify cilia in confocal z-stack images and determine their orientation based on the anatomical definition of the cilia base (broader) and tip (thinner) or the proximal-distal axis. The orientation angles were measured relative to the pia following the same approach established in (Yang et al., 2025), which were instead measured manually using ImageJ. Pericentrin staining confirmed that the broader end of cilia can always infer the location of the basal body. Although primary cilia become elongated in *reeler* mice, the proximal-distal morphology remains clearly distinguishable. The algorithm of Cilia Locator App uses a predefined multilayer pixel model of the ciliary nucleus to identify candidate by maximizing the correlation between the model and image intensity, followed by comparison with an orientation-dependent full-cilium model to estimate the most likely ciliary angle. Candidate locations and orientations (the proximal-distal axis) are visually presented to the user for confirmation, with options to adjust or reject detections if ambiguous, while accepted or rejected candidates are catalogued and excluded from subsequent searches to prevent duplicate identification. This iterative process continues until no additional cilia are detected, with previously identified cilia marked on subsequent images and an undo function available to correct user decisions. Collected cilia orientation data were exported into Excel files, which then were imported to another MATLAB® App, named Cilia Analysis (Mirhosseiniardakani et al., 2025). It was developed to analyze catalogued cilia orientations by fitting orientation data (0-360°) to one or two periodic normal distributions, with an optional uniform component to account for randomly oriented cilia. The analysis provided the peak orientation, standard deviation, proportion of cilia aligned with each major orientation, and goodness of fit based on the Kolmogorov-Smirnov (K-S) test. This App was also used to generate polar plots, histograms and color-coded spatial maps of cilia orientation.

Cilia orientation of early-born neurons was measured based on immunostaining of Ctip2 and AC3 antibodies. Cilia orientation of late-born neurons was measured based on immunostaining of Satb2 and AC3 antibodies. Cilia orientation of excitatory neurons was measured based on immunostaining of Glutaminase I and AC3 antibodies. AC3-NeuN co-staining generally staining neuronal primary cilia (from 80-95% excitatory neurons depending on location). GFAP and GAD65 antibodies were employed in immunostaining to differentiate GFAP-positive astrocytes and interneurons. GFAP-negative astrocytes may have missed the staining and cilia analysis, which is unlikely to change the conclusion.

### Color Coded Spatial Map of Cilia Orientation

To generate the color-coded spatial maps, confocal TIFF images were first uploaded into our custom MATLAB Cilia Locator app (Mirhosseiniardakani et al., 2025). Individual cilia were analyzed using this app, which calculated their orientation angles (5-degree resolution) and spatial coordinates and exported the measurements as an Excel file. The Excel file was subsequently imported into the custom MATLAB Cilia Analysis app, which automatically generated a spatial map. Each point represents one analyzed cilium at its corresponding x-y position within the original TIFF image, and its color represents the cilium orientation angle from 0° to 360°, as indicated by the color scale. The values on the x-and y-axes represent spatial coordinates relative to the coordinate system of the original TIFF image; they do not represent the direct distance of each cilium from the pial surface.

### Primary Cilia Length Measurement

Primary cilia length was measured manually in confocal z-stack images using the freehand-line tool in Fiji. Clearly distinguishable AC3-or Arl13b-positive cilia were selected, and a line was drawn along the ciliary axoneme from base to tip, following its curvature. The thicker end near the neuronal soma was identified as the base, whereas the thinner opposite end was identified as the tip (Sipos et al., 2018).

### Quadrant Analysis of Cilia Subcellular Location in Soma

NeuN and Glutaminase I immunostaining revealed the shapes of apical dendrites of principal neurons, which allowed cilia location identification. To evaluate the spatial distribution of primary cilia in various cell types and different brain regions, high-resolution confocal images were acquired using a 40X oil-immersion objective in Nikon’s Confocal microscope. Each soma/cell body was subdivided into four quadrants like apical, basal, left lateral, and right lateral by drawing perpendicular lines in Fiji (ImageJ) intersecting at the center of the nucleus, aligned with the apical-basal axis of the neuron. For interneurons and GFAP-positive astrocytes which often exhibit highly branched morphologies and do not have a single dominant primary process, we separated each cell body into four quadrants (top, bottom, left lateral, and right lateral). Each cilium base position was determined based on the quadrant in which the majority of the ciliary axoneme was localized. For each location, a minimum of 300 cilia per condition out of at least 3 animals were analyzed. The percentage of cilia in each quadrant was calculated and compared to determine whether they were equally distributed among the four orientations.

### Data Analysis and Statistics

Data analysis was conducted using Fiji, GraphPad Prism and Figure Panels were presented using CorelDRAW. Cilia orientation data were fit to several different distribution models in combination (uniform distribution - irregular cilia orientation; mono periodic normal distribution - one major cilia orientation; and double periodic normal distribution - two major cilia orientations). The Kolmogorov - Smirnov tests were applied to test the hypothesis that sampled orientation distribution could have been drawn from the fit probability distribution and facilitated the plotting of both the sampled orientation distribution and fit probability distribution (Mirhosseiniardakani et al., 2025). General statistical analyses were performed using ANOVA Tests or Kruskal-Wallis Tests for multiple group comparison. independent two-tailed Welch’s t-tests or Mann-Whitney U test to evaluate statistical significance, with a significance threshold of p < 0.05. Unless otherwise specified in the text or figure legends, significance levels are indicated as follows: ns (not significant); * p < 0.05; ** p < 0.01; *** p < 0.001. For the cilia-length analyses, each data point represents the length of an individual cilium. For the analyses of cilia location in somatic quadrants, each data point represents the percentage of cilia localized in the indicated quadrant for one mouse. The boxplots show the median and interquartile range, with individual data points displayed. Data were collected from 3-8 confocal images each animal and pooled from at least three mice of mixed sex in each age/genotype group. Minimal 100 neurons/cilia were analyzed from each mouse. A supplemental table (Table S2) provides detailed statistical results.

## Results

### The Primary Cilia of Pyramidal Neurons in Inside-out Laminated Regions are preferentially Localized to the Base of the Apical Dendrites and Orient toward the Pia

We assessed the orientation and subcellular location of primary cilia in the mouse cerebral cortex and other brain regions at P30. We first examined primary cilia in pyramidal neurons in the neocortex. Co-immunostaining for the neuronal marker NeuN, the basal body marker Pericentrin (Garcia and Reiter, 2016; Muhlhans et al., 2011), and the neuronal cilia marker AC3 (Bishop et al., 2007; Chen et al., 2016; Zhou et al., 2019) showed that AC3-positive primary cilia consistently originated from Pericentrin-stained basal bodies (Fig. 1A). Glutaminase I (a marker for excitatory neurons) staining further illustrated the localization of primary cilia in the somata of excitatory neurons (Fig. 1B). Cilia subcellular localization was classified based on the location of the cilia base within the soma (inferred from the broader end of the axoneme or the basal body staining). The soma was manually divided into four quadrants: apical, basal, lateral-right, and lateral-left in reference to the center of nucleus and the apical dendrite (Fig. 1C). We discovered that the majority of cilia were localized to the apical quadrant of somata, whereas substantially smaller proportions were found in the basal or lateral quadrants (Fig. 1C). Beyond the neocortex, we found that primary cilia are generally predominantly localized at the base of the apical dendrite (the apical quadrant) of principal neurons in inside-out laminated brain regions, which include entorhinal cortex, cingulate cortex, hippocampal CA1, and piriform cortex (Table S3).

**Fig. 1.**
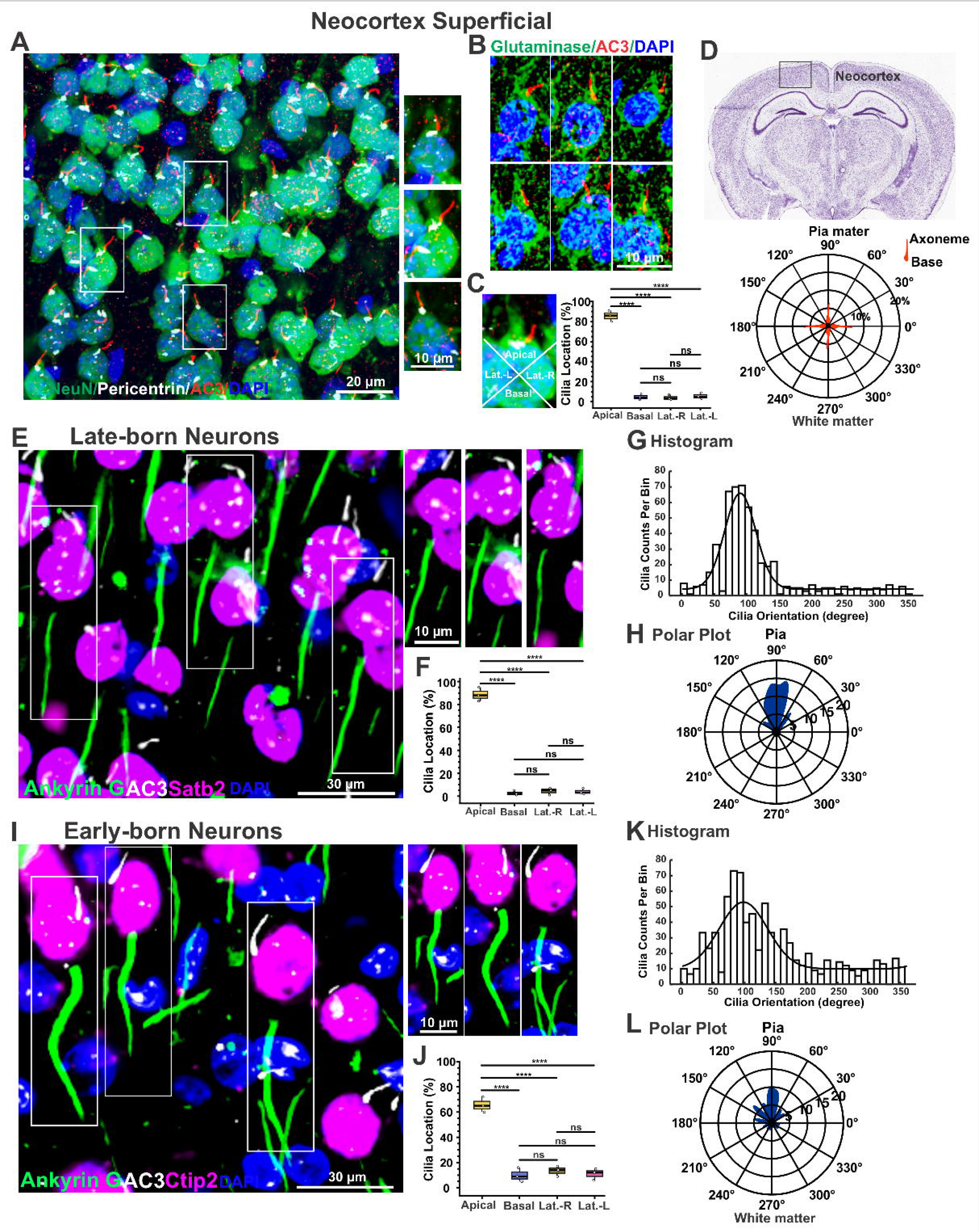
Primary cilia of neocortical principal neurons are predominantly localized to the base of apical dendrites and aligned opposite (and distal) to AIS. **(A-B)** Representative images showing cilia orientation and subcellular location in neurons of the superficial layers of P30. NeuN, AC3, and Pericentrin antibodies (A) labeled neurons, neuronal primary cilia and basal bodies, and Glutaminase I antibodies **(B)** selectively marked excitatory neurons. (C) Left, a diagram showing how four somatic quadrants (Apical, Basal, Lateral-Right and Lateral-Left) were defined in reference to the center of nucleus and the base of apical dendrite of pyramidal neurons. Right, primary cilia predominantly localize to the apical quadrant of neurons in the superficial layers. one-way ANOVA test. (D) An Allen brain atlas showing the imaging location and a polar plot indicating how cilia orientation was defined in reference to the pia mater and white matter. Pia-directed cilia orientation is 90 degrees. (E-H) Cilia location and orientation in late-born neurons marked by Satb2. (E) Representative images showing cilia of late-born neurons (marked by Satb2) aligned distally and opposite to AIS separated by the nuclei. Cilia and AIS were marked by AC3 and Ankyrin G, respectively. (F) Cilia location in 4 quadrants of late-born neurons. One-way ANOVA test. Cilia orientation histogram **(G)** and polar plot (H) of late-born neurons. (I-L) Cilia location and orientation in early-born neurons. (I) Representative images of cilia and AIS in early-born neuron marked by Ctip2. Cilia and AIS were marked by AC3 and Ankyrin G, respectively. **(J**) Cilia location in 4 quadrants of early-born neurons. Cilia of early-born neurons were mostly distributed to the apical quadrant, while some dispersed to other quadrants. One-way ANOVA test. Histogram (K) and polar plot (L) showing cilia orientation of early-born neurons. Histograms were fitted with one or two periodic normal distribution density functions, combined with a uniform distribution.

We next examined whether this subcellular location and directional orientation differed between late-born and early-born pyramidal neurons in the neocortex (Fig. S1). In late-born, Satb2-positive neurons, primary cilia showed a strong location preference for the apical quadrant (Fig. 1E, F). The histogram of cilia orientation displayed a narrow major peak around 90 degrees (the direction toward the pia), and polar plot analysis similarly demonstrated a strong concentration of cilia toward the pia matter (Fig. 1G, H). Intriguingly, the apical location of primary cilia is often accompanied with their orientation predominantly toward the pia, aligning distally and opposite to AIS at the basal side (Fig. 1E) (Fig. S1). In contrast, early-born, Ctip2-positive pyramidal neurons exhibited a bit broader distribution of cilia orientations (Fig. 1I-L) (Fig. S1). Although apically positioned cilia remained the predominant population, the proportion of apical cilia was lower, with increased presence in lateral and basal quadrants (Fig. 1J) (Fig. S1). The angular distribution of histogram of cilia orientation was substantially broader than that observed in late-born neurons, and the polar plot likewise showed a weaker pia-directed bias (Fig. 1K, L). These results demonstrate that primary cilia of neocortical pyramidal neurons preferentially localize to the apical side of the soma and orient toward the pia. This directional organization is particularly prominent in late-born neurons, whereas early-born neurons display a more heterogeneous cilia location and orientation.

Atypical pyramidal neurons in the deep neocortex, which deviate from the classical morphological and electrophysiological features of pyramidal neurons (Landrieu and Goffinet, 1981), and some have inverted, while others have oblique apical dendrites (Steger et al., 2018; Steger et al., 2013). Immunostaining for AC3 and Glutaminase I (or NeuN) revealed abundant cilia in these cells (Fig. S2A-B). However, primary cilia do not align distally or opposite to AIS in reference to the nuclei, and the relative position of cilia and AIS appears irregular (Fig. S2C). The histogram and polar plot of cilia orientation angles demonstrated a broad, evenly dispersed distribution without a dominant peak (Fig. S2D), indicating a lack of preference for directional orientation. The analysis of centriole/cilia subcellular location also revealed broad distribution among four somatic quadrants, indicating the absence of location bias (Fig. S2E). Hence, unlike pyramidal neurons, atypical pyramidal neurons in the deep neocortical layers do not exhibit directional cilia orientation or preference for apical location.

Pyramidal or excitatory neurons in the amygdala are non-laminated but organized into distinct nuclei (LeDoux, 2007). We found that their cilia pattern somewhat resembled that of atypical pyramidal neurons in the deep neocortex: primary cilia do not align distally or opposite to AIS, show preferred orientation, or preferably localize to the base of the apical dendrites (the apical quadrant) (Fig. S3A-D). Consistently, in other non-laminated brain regions examined, such as the hypothalamus, striatum, and thalamus, neuronal primary cilia were broadly positioned in the soma, lacking directionality (Table S3). Additionally, the primary cilia of interneurons and GFAP-positive astrocytes throughout the cerebral cortex were also distributed broadly around the nucleus (Fig. S3E, F) and lack a directional orientation or preference for somatic location (Table S3).

Together, the primary cilia of pyramidal neurons in inside-out laminated brain regions preferentially localize at the base of apical dendrites (the apical quadrant) and predominantly orient toward the pia direction, while atypical pyramidal neurons, excitatory neurons in non-laminated regions, interneurons and GFAP-positive astrocytes do not exhibit that pattern.

### Reelin Deficiency Disrupts Pia-Directed Orientation of Cilia

We asked what factor(s) regulate the cilia orientation of cortical principal neurons. Reelin is a key extracellular cue that influences neuronal placement, apical-basal neuronal polarity, and laminar formation in the cerebral cortex (Jossin, 2020; Rice and Curran, 2001). We hypothesized that Reelin regulates the subcellular localization and directional orientation of primary cilia of cortical principal neurons. To test this, we compared cilia orientation and cilia subcellular location first in the superficial neocortical layers using *reeler* and WT mice. *reeler* carries a mutation in the Reelin gene (*Reln*) and is an extensively characterized null model (D’Arcangelo et al., 1995). We found that primary cilia in WT mice exhibited a predominant orientation toward the pia direction (Fig. 2A, B). Cilia orientation (Fig. 2A-D left) showed a strong peak around 90°, with most cilia oriented toward the pia. In contrast, *reeler* superficial layer neurons showed diverse cilia orientation, lacking predominant alignment toward the pial surface (Fig. 2A-B). The corresponding histogram and polar plot of cilia orientation revealed a more dispersed distribution of cilia orientation angles. Data fitting indicated that WT cilia orientation followed a single periodic normal distribution centered around 90°, while *reeler* cilia mostly presented a more uniform distribution, indicative of randomized cilia orientation (Fig. 2C, D). Color-coded spatial maps (Fig. 2E) of pooled cilia orientation data further supported this observation: WT samples showed a pattern with major cilia orientation, while *reeler* samples exhibited a more heterogeneous and disorganized orientation in the superficial layer.

**Fig. 2.**
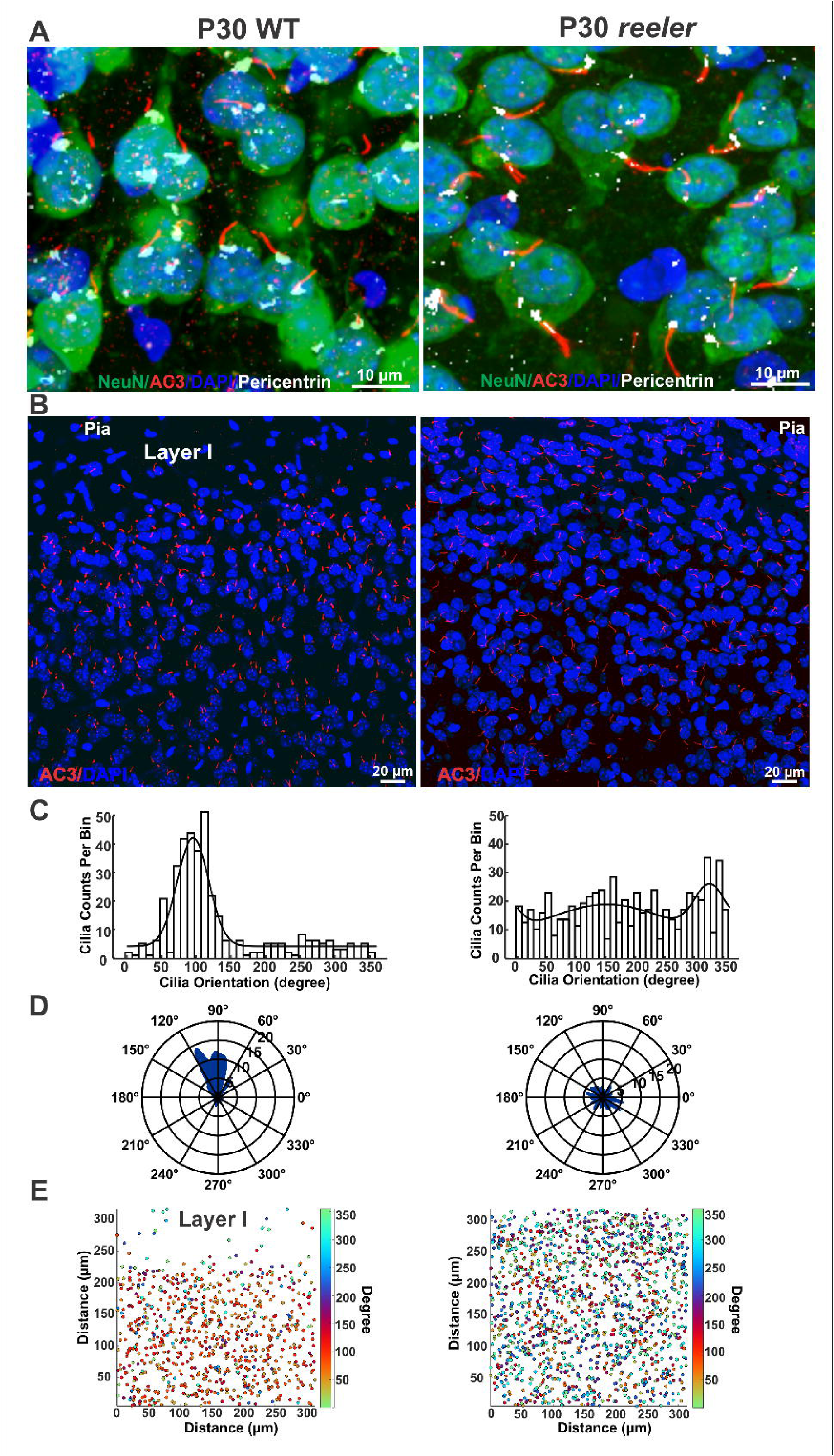
Disrupted directional cilia orientation in the superficial layer of the *reeler* neocortex. **(A-B)** Primary cilia in the superficial layers of the WT and *reeler* neocortices. (**A**) Co-immunostaining of NeuN, Pericentrin and AC3. WT cilia were mostly localized to the apical dendrites and oriented to the pia, while *reeler* cilia were broadly distributed in the soma and lacked preferred orientation. **(B**) Representative large-view images showing orientation of AC3-labeled neuronal cilia in WT and *reeler*. **(C-D)** Histograms and polar plots of cilia orientations in the superficial layers of the WT and *reeler* neocortices. WT cilia are predominantly oriented toward the pia (peak at ∼90°), while *reeler* cilia do not. Data were fitted with one or two periodic normal distribution density functions, combined with a uniform distribution. **(E)** Color-coded spatial map of cilia orientation in the superficial layers visualizing cilia orientation across the sampled fields. Data from 354 WT cilia (from 3 mice) and 557 *reeler* cilia (from 3 mice) were pooled to construct each spatial map. Note that WT has no excitatory neurons in Layer I, while *reeler* does not have distinguishable Layer I.

### Reelin Deficiency Partially Alters the Apical Localization and Pia-directed Orientation of Primary Cilia in Late-Born Pyramidal Neurons

Nevertheless, *reeler* is known to exhibit “outside-in” or “inverted” cortical lamination pattern and the superficial layer of *reeler* neocortex mostly contains early-born neurons, while its deep layer accommodates late-born neurons (D’Arcangelo et al., 1995; Ogawa et al., 1995), although the layer-specific boundary of *reeler* is not as sharp as WT (Hoffarth et al., 1995). To distinguish early-born from late-born neurons, we used Ctip2 antibody and Satb2 antibody to label early-and late-born neurons, respectively.

The cilia pattern in cortical principal neurons, including directional orientation, are subject to changes during postnatal development (Yang et al., 2025). We therefore examined the developmental changes in the subcellular localization and orientation of primary cilia in late-born neurons from P5 to P30 (Fig. 3). In WT mice, primary cilia of late-born neurons were already preferentially localized to the apical quadrant of neuronal somata at P5, with substantially fewer cilia located in the basal or lateral quadrants (Fig. 3A-C). This preferential apical localization remained prominent during postnatal development and was maintained through P30 (Fig. 3D). Cilia orientation, however, underwent a marked progression with postnatal development (Fig. 3E). At P5, cilia orientation in WT late-born neurons was relatively broadly distributed, without a prominent directional peak. By P10, however, a distinct peak emerged around the pia-directed angle, and this directional distribution became increasingly prominent at P14 and P30 (Fig. 3E). Therefore, the preferential apical localization of primary cilia appears earlier than their directional orientation, whereas pia-directed cilia orientation is progressively established during postnatal development.

**Fig. 3.**
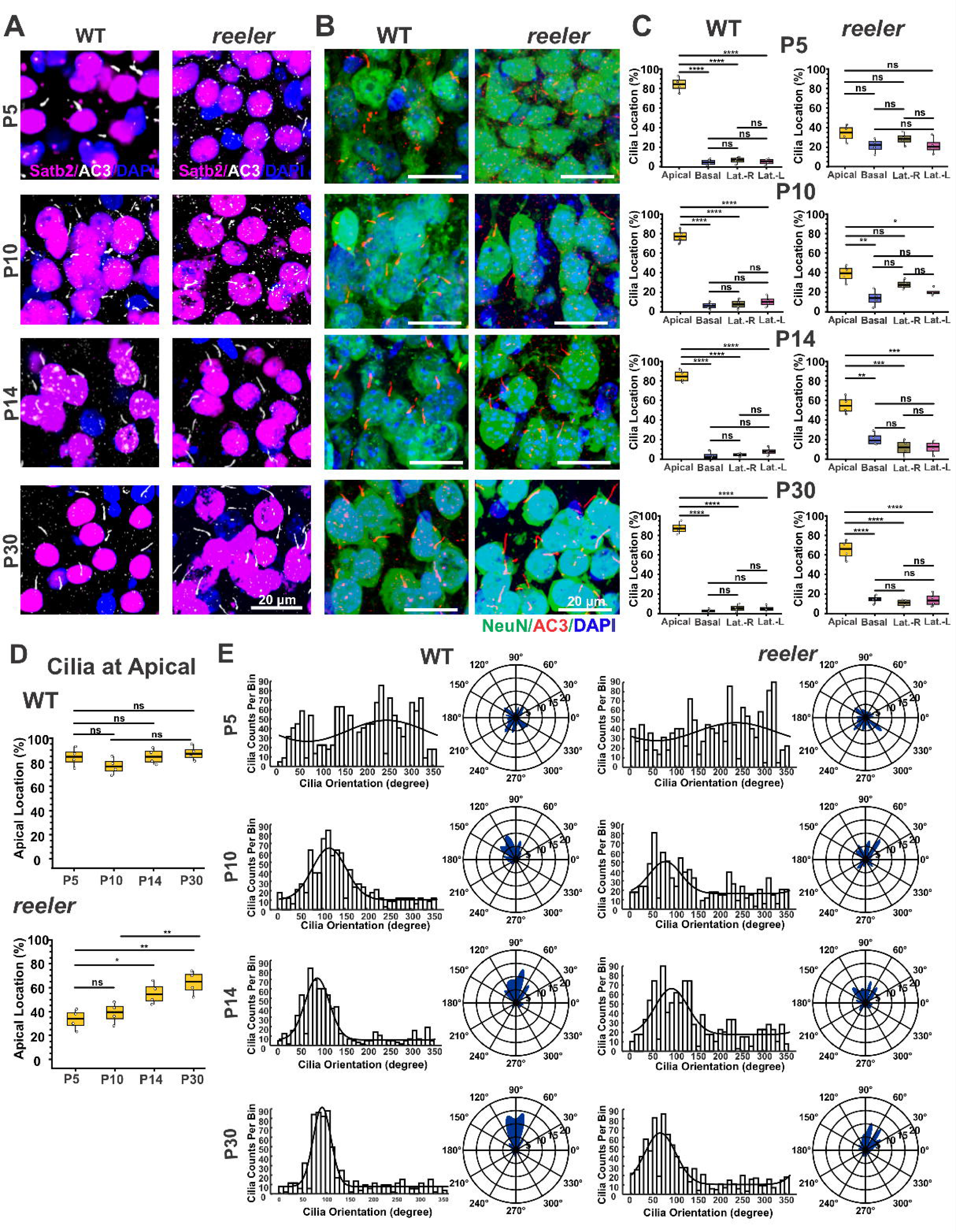
*Reeler* late-born neocortical neurons exhibit a postnatal correction in cilia apical localization and directional orientation. **(A-B)** Representative images of WT and *reeler* late-born neurons and primary cilia at P5, P10, P14 and P30. Primary cilia, AIS, late-born neurons were labeled by AC3 and Satb2 (**A**) or NeuN (**B**) antibodies, respectively. (**C**) Cilia location in four quadrants of late-born neurons of WT and *reeler* from P5 to P30. One-way ANOVA tests were used. (**D**) The changes of cilia location in apical quadrant late-born neurons from P5 to P30 of WT vs Reeler. *reeler* cilia exhibit increased presence in the apical quadrant from P5 to P30, while WT cilia remained strong presence. (**E**) Cilia orientation changes in histograms and polar plots from P5 to P30 of WT and *reeler*.

In *reeler* mice, the developmental process of primary cilia of late-born neurons was markedly altered (Fig. 3). At P5, cilia were localized more broadly among the apical, basal, and lateral quadrants compared with WT neurons, indicating that preferential apical localization was impaired at this early postnatal stage. At P10-P30, *reeler* late-neurons continued to exhibit a more heterogeneous distribution of cilia in the soma, although an apical bias was detectable, particularly at later ages (Fig. 3C). Reelin deficiency substantially reduces the preferential localization of primary cilia to the apical quadrant of late-born pyramidal neurons at early age; however, at later stages, a greater proportion of cilia becomes distributed within the apical quadrant (Fig. 3D).

Cilia orientation was also altered in *reeler* mice. Whereas WT neurons progressively developed a prominent pia-directed peak between P10 and P30, *reeler* neurons displayed a broader and less sharply defined orientation distribution. Nevertheless, a directional bias became evident in *reeler* neurons at later postnatal ages, suggesting that a subset of neurons partially establish directional cilia orientation despite the absence of Reelin (Fig. 3E). These findings indicate that Reelin deficiency partially alters the preferential apical localization and directional orientation of primary cilia in late-born neurons.

### Reelin Deficiency Completely Disrupts Apical Cilia Localization and Directional Cilia Orientation of Early-born Neurons

We next examined cilia phenotypes in early-born pyramidal neurons to evaluate whether the effects of Reelin deficiency differ in neuronal populations (Fig. 4A-B). In WT mice, early-born neurons showed a developmental progression in ciliation pattern. At P5, primary cilia were preferentially localized to the apical quadrant, although their orientation was relatively heterogeneous. By P30, the primary cilia of WT early-born neurons maintained strong preferential apical localization and developed a prominent directional orientation, with the majority of cilia oriented toward the pia (Fig. 4C-G). In contrast, *reeler* early-born neurons failed to develop either of these features. At both P5 and P30, primary cilia were distributed relatively evenly among the apical, basal, and lateral quadrants, with no significant preference for the apical quadrant (Fig. 4C-E). Consistent with the lack of preferential subcellular localization, cilia orientation remained broadly distributed from P5 to P30, rather than developing the prominent directional peak (Fig. 4F-G). These results reveal a particularly strong effect of Reelin deficiency on primary cilia of early-born neurons.

**Fig. 4.**
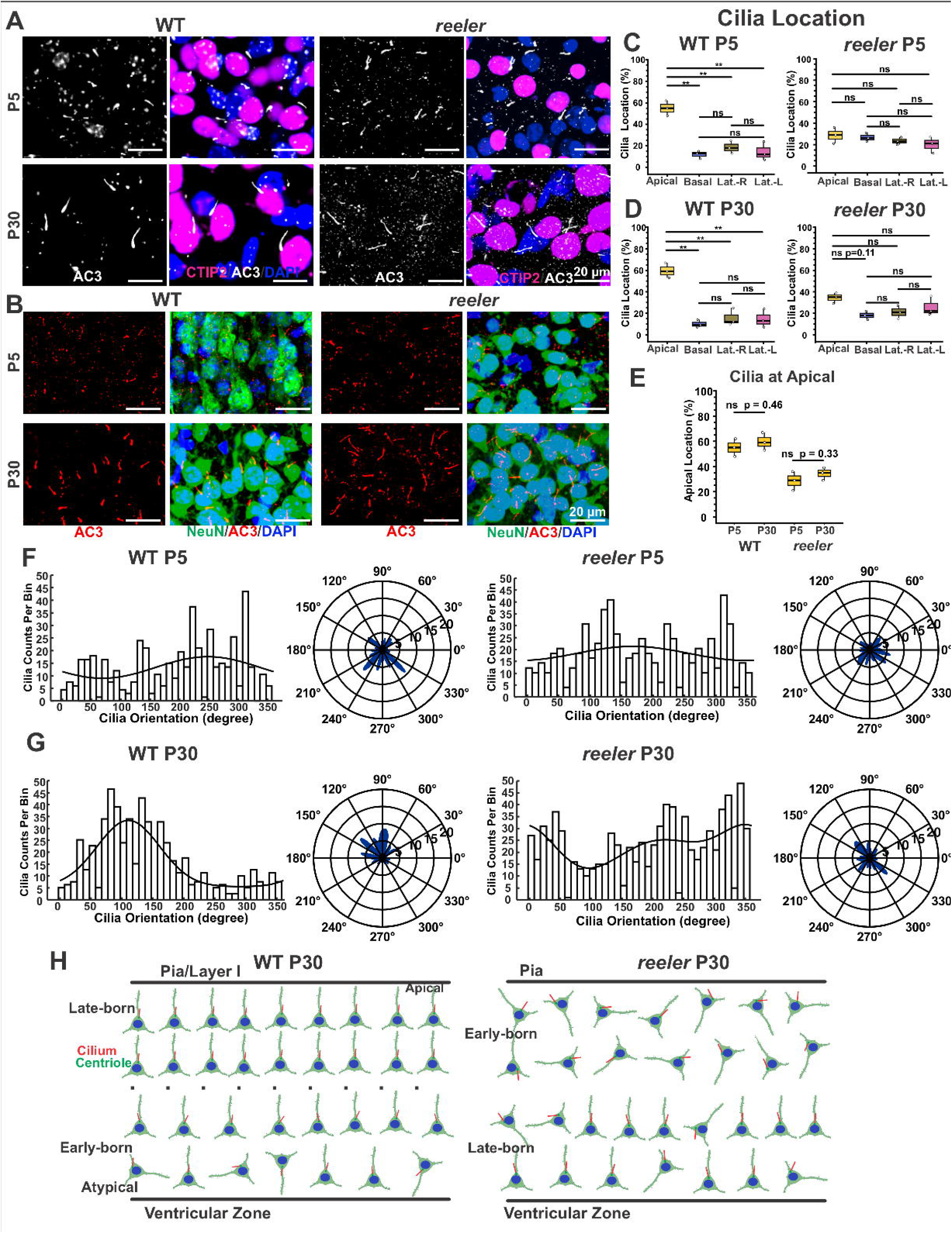
*reeler* early-born neocortical neurons do not undergo postnatal correction in their cilia orientation and apical location. **(A-B)** Representative images of primary cilia of WT and *reeler* early-born neurons at P5 and P30 marked by Ctip2 and NeuN, respectively. (**C-D**) Cilia subcellular location in four quadrants in WT and *reeler* early-born neurons of P5 and P30. (**E**) Changes in the percentage of cilia in the apical quadrant in early-born neurons of WT and *reeler*, P5 vs P30. (**F-G**) Histograms and plot plots of cilia orientation of WT and *reeler* early-born neurons at P5 and P30. WT cilia become more oriented toward the pia direction at P30, while *reeler* exhibit few change in cilia orientation from P5 to P30. (**H**) Schematic diagrams depicting cilia orientation and location in early-and late-born neocortical neurons of WT and *reeler* of P30.

The partial restoration of apical localization and directional orientation in *reeler* late-born neurons at later postnatal stages suggests that late-born neurons retain a greater capacity to correct mis-orientation in the absence of Reelin. In comparison, *reeler* early-born neurons apparently lack this feature (Fig. S4). The scheme in Fig. 4H summarizes this distinction, showing disrupted cilia orientation and location in misplaced early-born *reeler* neurons but comparatively stronger apical and directional organization among late-born neurons.

### Reelin Deficiency Partially Alters the Apical Localization and Directional Orientation of Primary Cilia in CA1 Pyramidal Neurons

We next examined the impacts of Reelin deficiency on primary cilia in the hippocampal CA1 region, where the developmental changes in cilia phenotypes in WT pyramidal neurons have been characterized previously (Yang et al., 2025). In the WT CA1 region, pyramidal neurons, most of which exhibit late-born features, are organized into superficial and deep populations within the stratum pyramidale (SP) (Cembrowski and Spruston, 2019). AIS marked by Ankyrin G is typically identified in the basal side (Pan et al., 2006). Primary cilia in CA1 SP exhibit two approximately opposite directional orientations toward the stratum radiatum (SR) and stratum oriens (SO), respectively (Yang et al., 2025). CA1 neuronal primary cilia emerge after P7 (Yang et al., 2025). Their major cilia orientation varied slightly with postnatal age (Fig. 5A-F), consistent with our previous report (Yang et al., 2025). In *reeler* mice, the layering of the CA1 pyramidal layer was disrupted (Landrieu and Goffinet, 1981; Ogawa et al., 1995; Rice and Curran, 2001), and neurons were dispersed and roughly separated into two layers (Fig. 5A). We therefore divided *reeler* CA1 neurons into top and bottom populations according to their relative positions within the dispersed neuronal layer. Despite the severe disruption of neuronal placement, cilia orientation was not completely randomized in *reeler* (Fig. 5C). At P10 and P14, *reeler* neurons in the top population exhibited a prominent directional peak, whereas the bottom population showed a broader orientation distribution. By P30, directional organization remained apparent in the top population, while the bottom population developed a more evident bidirectional pattern (Fig. 5D-F). These results indicate that Reelin deficiency does not prevent CA1 pyramidal neurons from establishing directional cilia orientation.

**Fig. 5.**
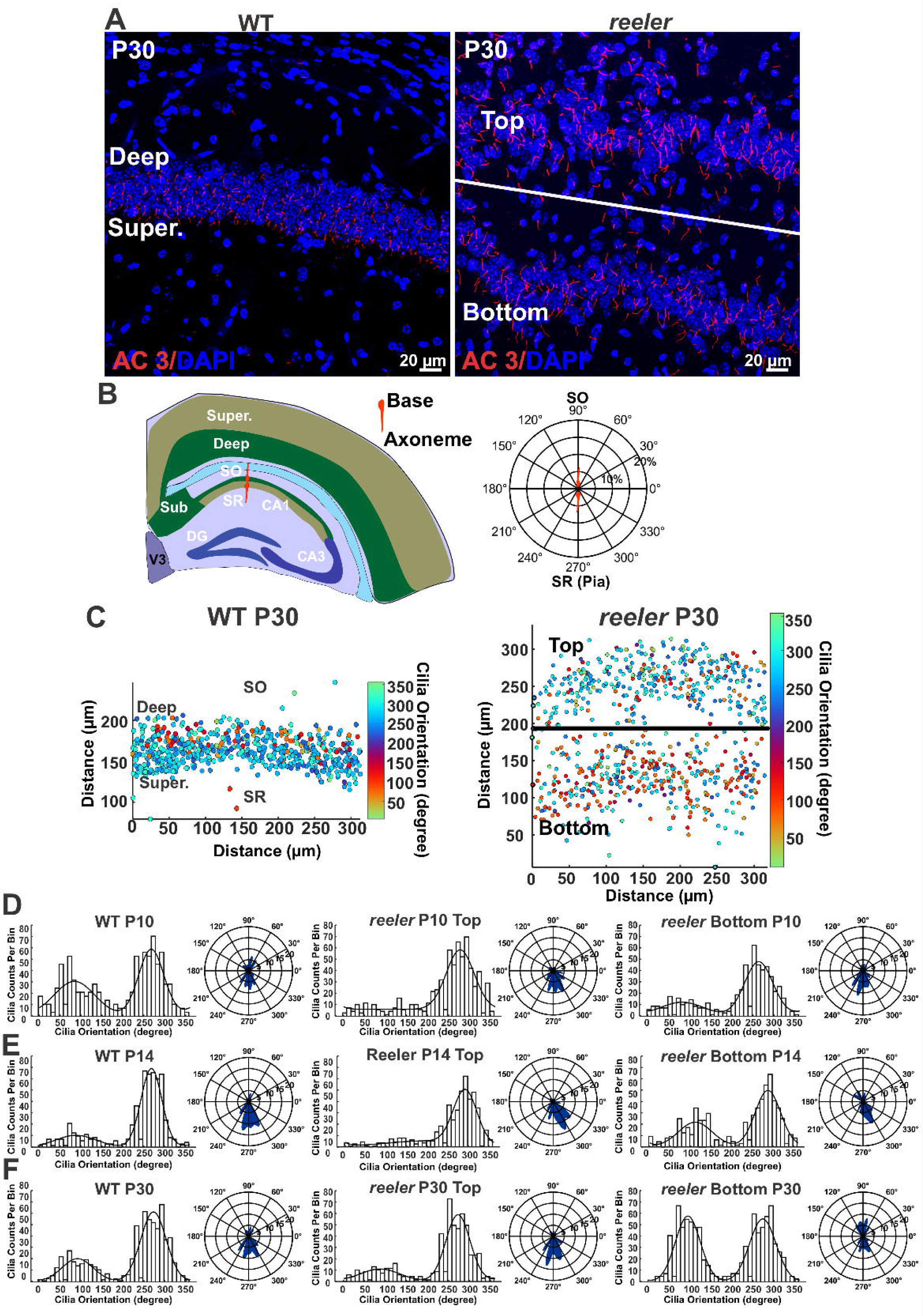
Reelin deficiency partially affects the orientation of primary cilia in CA1 pyramidal neurons. **(A)** Representative cilia images in WT and *reeler* CA1 region of P30. *reeler* CA1 stratum pyramidale was split into “top” and “bottom” sublayers. (**B**) Cilia orientation measurement in references to CA1 anatomy and polar plot. **(C)** Spatial map of cilia orientation in WT and *reeler* CA1, visualizing cilia orientation across the sampled fields, color-coded by angle (degrees). **(D-F)** Histograms of cilia orientation in the WT CA1 of P10, P14 and P30. Histograms of cilia orientation in the top and bottom sublayers of *reeler* at P10 (**D**), P14 (**E**), and P30 (**F**).

We further examined whether Reelin deficiency affects the subcellular localization of primary cilia (Fig. 6). In WT CA1 pyramidal neurons, primary cilia were preferentially localized to the apical quadrant of the soma at P10, and this apical preference remains prominent at P30 (Fig. 6A-D). In contrast, cilia in *reeler* CA1 neurons at P10 were more broadly distributed among the apical, basal, and lateral quadrants, lacking predominant preference for the apical quadrant. Thus, similar to the phenotype observed in the *reeler* neocortex, Reelin deficiency disrupts the preference for apical localization of primary cilia at early stage. However, the cilia localization in *reeler* CA1 was substantially corrected during postnatal development. By P30, *reeler* primary cilia again showed a strong preference for the apical location, despite persistent neuronal misplacement (Fig. 6A-D). Together with the developmental changes in cilia orientation shown in Fig. 5, these results suggest that primary cilia in *reeler* CA1 pyramidal neurons retain the capacity to partially restore apical localization and directional orientation.

**Fig. 6.**
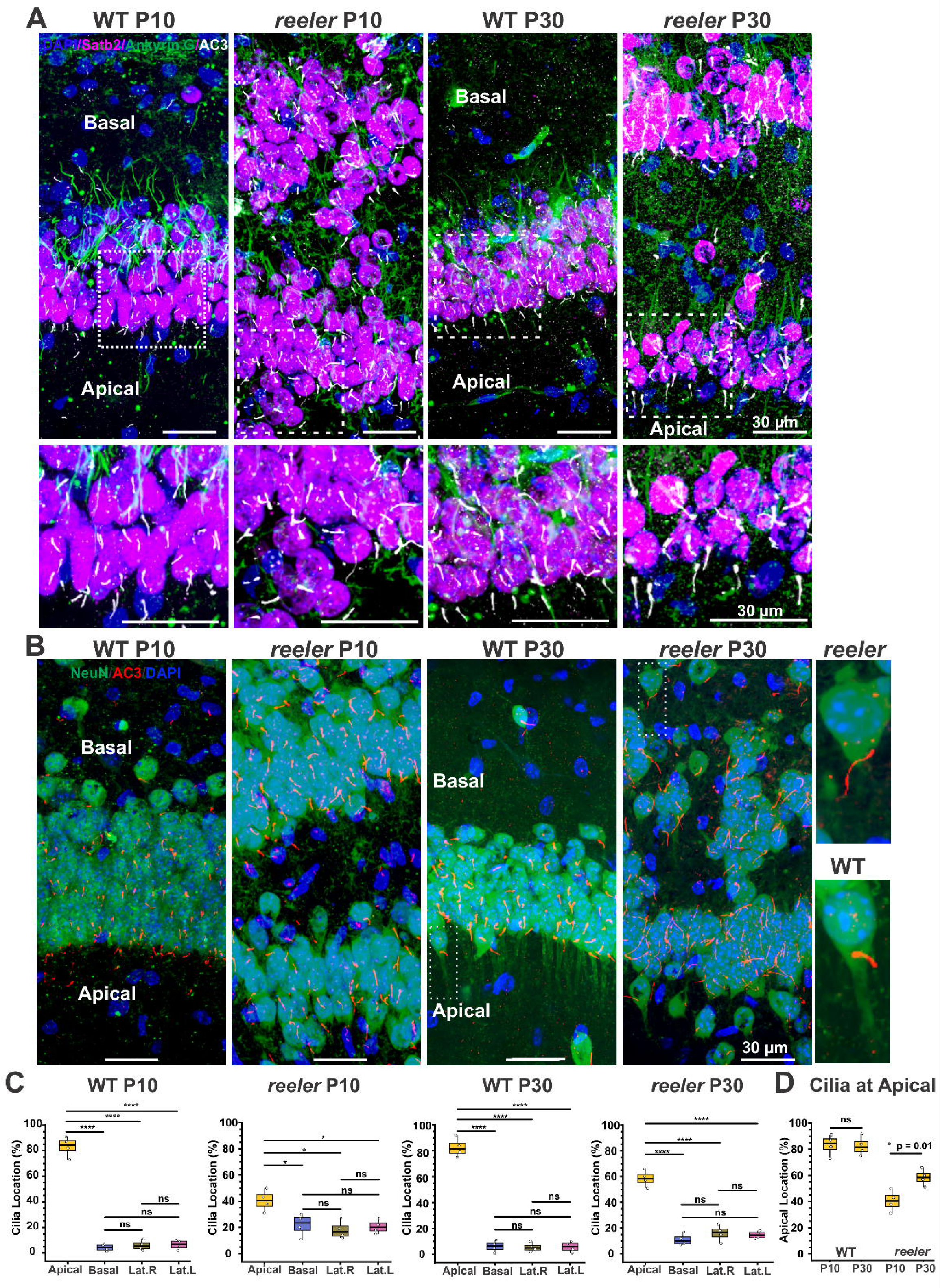
Reelin deficiency partially alters the apical localization of cilia in CA1 pyramidal neurons. (**A**) Representative images showing relative position of primary cilia (marked by AC3) and AIS (marked by Ankyrin G) in reference to nuclei in WT and *reeler* CA1 region of P10 and P30. White boxed regions were enlarged at the bottom. WT neurons typically have cilia in the apical side and AIS in the basal side at both P10 and P30. *Reeler’s* cilia-at-apical pattern was impaired at P10 but partially corrected at P30. (**B**) NeuN staining roughly visualizes pyramidal cell shape and relative location of cilia in WT and *reeler*. Right insets are enlarged neuron examples of P30. **(C)** Cilia location in four quadrants of WT and *reeler* neurons at P10 and at P30. One-way ANOVA tests. (**D**) The change of cilia location in the apical quadrant of CA1 neurons, WT and *reeler* of P10 and P30. Student’s T-tests were used.

### Reelin Deficiency Disrupts the Orientation and Apical Location of Primary Cilia in Evolutionarily Conserved Cortical Regions

Next, we examined cilia phenotypes in the hippocampal DG and CA3, which are considered highly conserved allocortical structures (Blackstad et al., 2022; Krubitzer and Seelke, 2012), compared to the CA1 and neocortex. In WT DG of P30, primary cilia exhibited a preference for the apical quadrant of the soma, with fewer cilia localized to the basal or lateral quadrants (Fig. 7A-D). Cilia orientation displayed a bidirectional pattern, with two major orientation peaks separated by approximately 180° (Fig. 7E), in line with a previous report (Yang et al., 2025). In *reeler* DG of P30, both features were markedly disrupted. Primary cilia were more broadly distributed among the apical, basal, and lateral quadrants, and the apical location preference was no longer evident (Fig. 7A-D). Consistent with this altered subcellular localization, the orientation peaks observed in WT DG were absent in *reeler*, resulting in a substantially broader and less directional preference of cilia orientations (Fig. 7E).

**Fig. 7.**
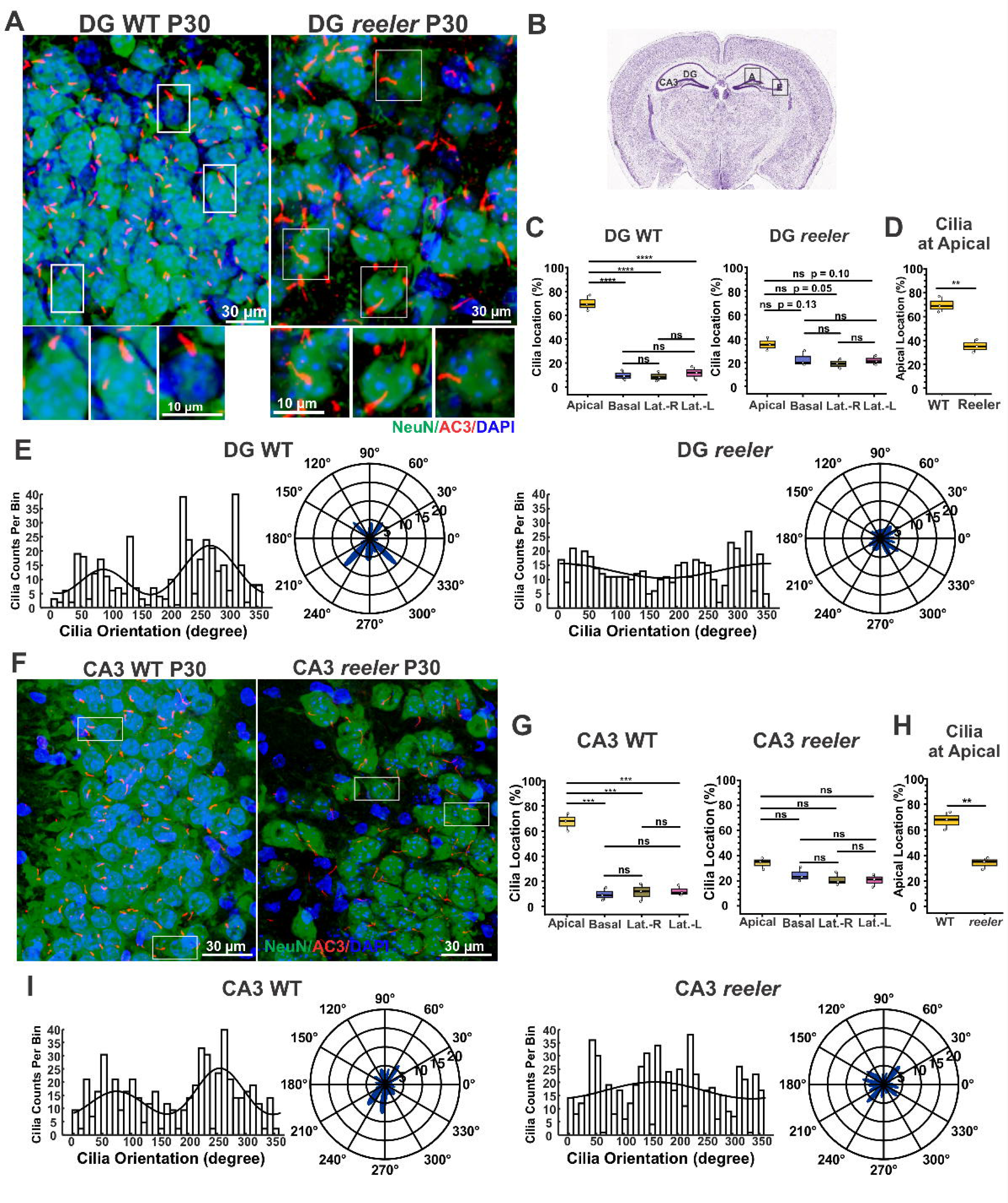
Disrupted cilia subcellular location and orientation in the *reeler* DG and CA3 of P30. **(A)** Representative neurons and cilia images in WT and *reeler* DG of P30, marked by NeuN and AC3 antibodies, respectively. Images were acquired from the suprapyramidal blade of the dentate gyrus. (**B**) An Allen brain atlas showing the imaging location in DG (and CA3 as well). **(C)** Cilia subcellular location in WT and *reeler* DG neurons. In WT, cilia location showed a strong preference to the apical quadrant. In contrast, *reeler* cilia displayed no location preference to any quadrant, one-way ANOVA test. (**D**) Cilia location in the apical quadrant DG neurons, WT vs *reeler*. (**E**) Histograms and polar plots of cilia orientation in the WT and *reeler* DG of P30. WT cilia roughly had opposite cilia orientations, while *reeler* do not. **(F)** Representative cilia images in the WT and *reeler* CA3 of P30. (**G**) Cilia location in WT and *reeler* CA3 neurons. In WTs, cilia were predominantly localized to the apical quadrant. In contrast, *reeler* CA3 cilia had no preference to 4 locations. (**H**) Cilia localization at the apical quadrant CA3 neurons, WT vs *reeler.* (**I**) Histograms and polar plots of cilia orientation in WT and *reeler* CA3 neurons of P30.

A similar phenotype was observed in CA3. In WT CA3 of P30, primary cilia were preferentially localized to the apical side of neuronal somata, with the apical fraction significantly greater than the basal and lateral fractions (Fig. 7F-H). Cilia also exhibited a directional pattern, characterized by two major orientation peaks rather than an even distribution (Fig. 7I). In contrast, cilia in *reeler* CA3 were distributed more evenly among the four somatic quadrants, with no significant preference for apical localization. Cilia orientation was likewise broadly distributed, and the prominent directional pattern observed in WT CA3 was largely absent (Fig. 7F-I).

The piriform cortex, a major part of the olfactory cortex, is 3-layered paleocortex and a more primitive cortical structure than the hippocampal CA1 and neocortex (Klingler, 2017). The piriform cortex overall forms in an inside-out developmental manner, albeit with a sublayer inversion in Layer II (Martin-Lopez et al., 2019). We examined cilia orientation and cilia subcellular location in Layer II and III of the WT piriform cortex of P30 (Fig. 8). In Layer II, which is a compact lamina, neuronal primary cilia exhibited two major opposite directions (Fig. 8C), while in Layer III, which is sparsely layered, cilia had one major direction, predominately oriented toward the pia direction (Fig. 8D). Consistently, most of cilia in the piriform cortex were localized at the base of apical dendrite (the apical quadrant) (Fig 8E). Reelin deficiency disrupted proper neuronal placement, laminated structures, and apical-basal neuron orientation in the piriform cortex (Fig. 8). Moreover, primary cilia in both superficial and deep layer of the *reeler* piriform cortex no longer exhibited preferred orientation toward the pia direction (Fig. 8F-G). Consistently, cilia were no longer confined to the apical quadrant but became broadly distributed to 4 somatic quadrants (Fig. 8H). The preferences of cilia location to the apical quadrant exhibit a strong difference between WT and *reeler* neurons (Fig. 8I). These data demonstrate that Reelin deficiency disrupts the directional orientation and subcellular location of neuronal primary cilia in the piriform cortex. The impaired phenotypes were maintained at P30, suggesting a lack of postnatal correction.

**Fig. 8.**
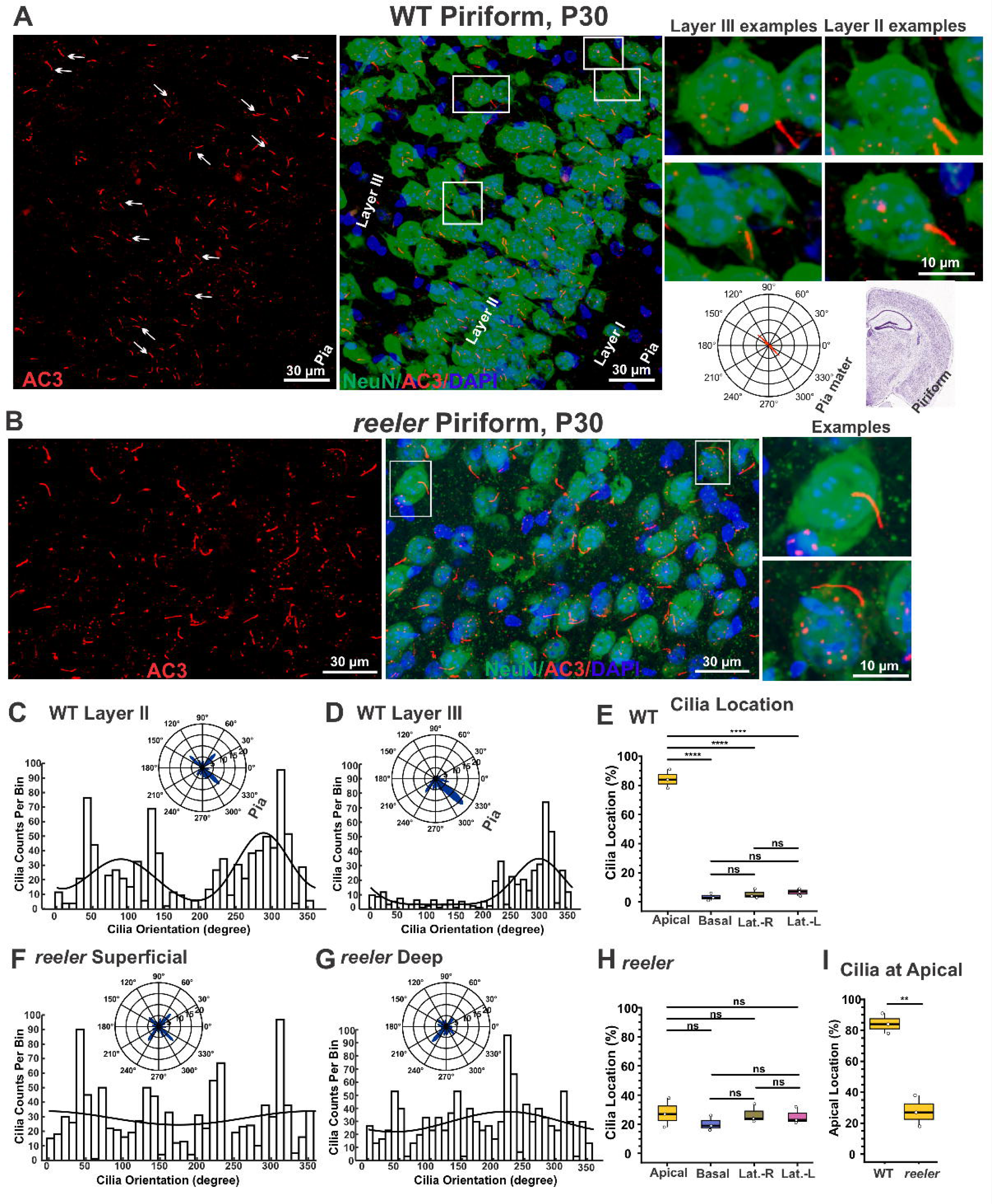
Disrupted cilia orientation and apical localization in the *reeler* piriform cortex of P30. Primary cilia in the WT **(A)** and *reeler* **(B)** piriform cortex of P30. Representative individual neuron images enlarged on the right. Inset polar plot and Allen brain atlas illustrating cilia orientation in the piriform cortex relative to the pia mater. **(C-E)** Histograms and polar plots of cilia orientation in Layer II (**C**) and III in the WT piriform cortex (**D**), and cilia location in 4 somatic quadrants of WTs (**E**). Histograms and polar plots of cilia orientation in superficial and deep layers of *reeler* piriform cortex (**F-G**), and cilia location in somatic quadrants in the *reeler* piriform cortex (**H**). One-way ANOVA tests were used to examine cilia location effects. (**I**) Percentage of cilia subcellular location to the apical quadrant of WT vs *reeler* at P30. Two-tailed unpaired Welch T-test.

Together, Reelin deficiency strongly impairs the directional orientation and apical localization of primary cilia in principal neurons in evolutionary conserved laminated regions, including the DG, CA3 and piriform cortex. These cortical regions lack strong postnatal corrections for re-orientation in *reeler* mice at least as of P30.

### Primary Cilia of Principal Neurons in the *Reeler* Cerebral Cortex Continue to Elongate at a Developmental Stage when WT Cilia Length Stabilize

Next, to examine if Reelin deficiency affects cilia length, we measured cilia length across different brain regions in WT and *reeler* mice. We performed staining using AC3 antibody to specifically mark neuronal primary cilia (Bishop et al., 2007; Chen et al., 2016; Zhou et al., 2019) and Arl13b antibody to label astrocytic primary cilia (Sterpka and Chen, 2018; Sterpka et al., 2020). We analyzed cilia in WT and *reeler* mice of P14 and P30, first focusing on laminated brain structures including the hippocampal CA1, CA3, DG, superficial and deep layers of the neocortex, and piriform cortex (Fig. 9).

**Fig. 9.**
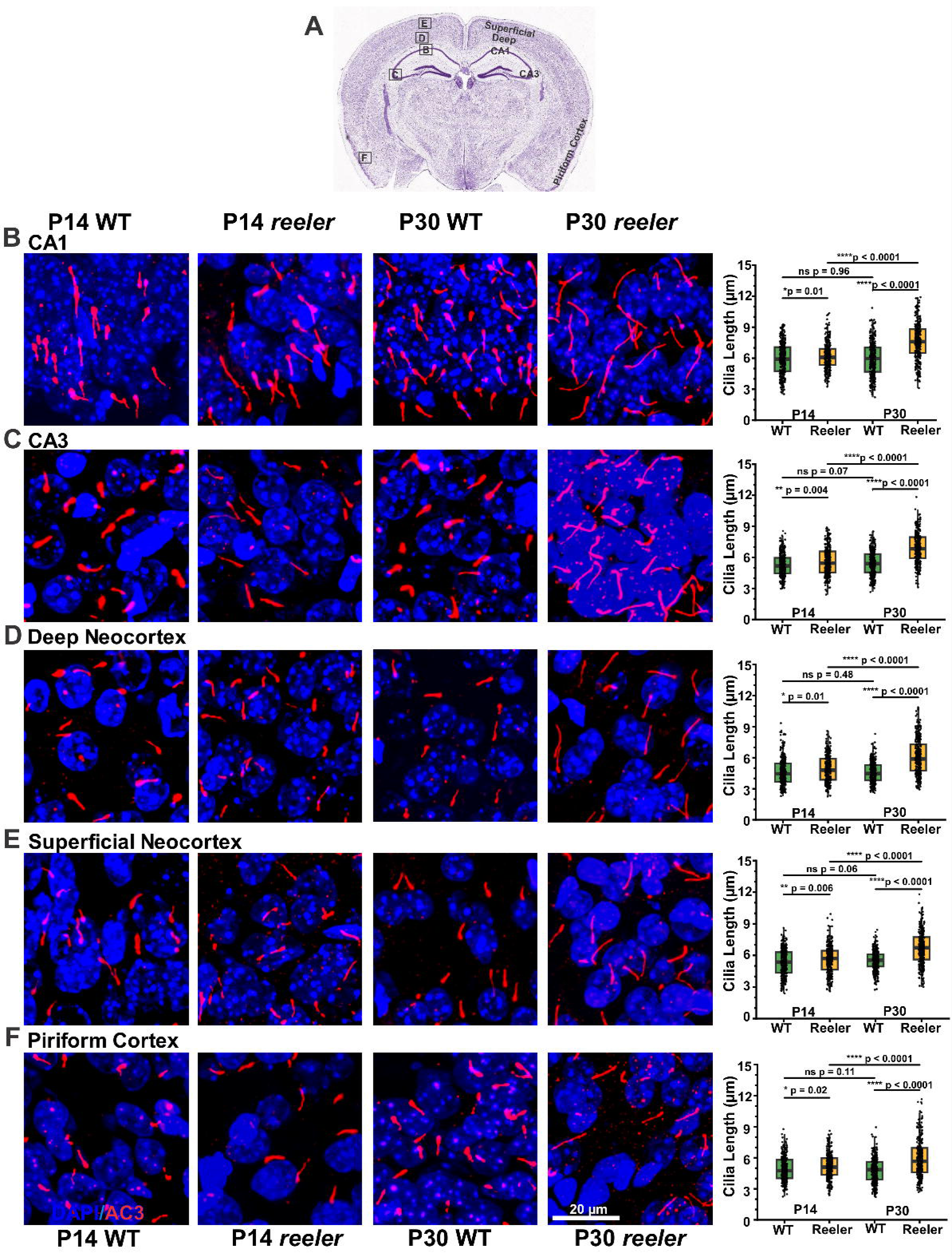
Reelin deficiency promotes cilia elongation after P14 in regions across the cerebral cortex. (**A**) Allen brain atlas indicate regions examined. **(B)** CA1; **(C)** CA3; **(D)** Deep Neocortex; **(E)** Superficial Neocortex; and **(F)** Piriform Cortex. Left, representative images of neuronal primary cilia (marked by AC3 antibody) in various cortical regions of WT and *Reeler* of P14 and P30. Right, scatter plots showing cilia length of P14 and P30 WT and *reeler* mice. Two-tailed unpaired Welch’s T-tests were performed to compare two groups.

In the hippocampal CA1 region, neuronal primary cilia were longer in *reeler* than in WT of P30 (Fig. 9B, left). Quantitative analysis of cilia length (Fig. 9B, right) confirmed a significant increase in cilia length in *reeler* mice. In fact, this trend of longer cilia was consistently observed in all other laminated regions analyzed, including CA3, DG, superficial neocortex, deep neocortex, piriform cortex, and entorhinal cortex (Fig. 9C-F, Table S4). Notably, at P14, cilia lengths in *reeler* mice were comparable to WT controls across all examined regions, with minor differences detected (Fig. 9B-F, P14 groups). Additionally, WT mice exhibited comparable cilia lengths between P14 and P30, consistent with previous report (Yang et al., 2025). In contrast, *reeler* mice showed a pronounced elongation of primary cilia from P14 to P30 in all laminated brain regions examined. This age-dependent effect suggests that Reelin directly or indirectly constrains cilia elongation during postnatal cortical development. Further, we found that Reelin haploinsufficiency do not significantly affect cilia length in principal neurons in regions across the cerebral cortex (Fig. S5), arguing against a dosage effect.

### Reelin Deficiency Does Not Affect the Cilia Length of Excitatory Neurons in Non-Laminated brain Regions, Interneurons or GFAP-positive Astrocytes

To determine whether Reelin influences cilia length in non-laminated brain regions, we measured cilia length in several non-laminated areas in WT and *reeler* mice of P30. These regions included the thalamus, striatum, hypothalamus, and amygdala (Fig. S6A). In the thalamus (Fig. S6B), immunofluorescence images (right) show similar cilia length between WT and *reeler* mice, and the box graph (left) confirms no significant difference. Likewise, in the striatum (Fig. S6C), hypothalamus (Fig. S6D), and amygdala (Fig. S6E), there were no differences in cilia length between WT and *reeler*. These findings indicate that Reelin deficiency does not affect cilia length in non-laminated regions of the brain.

To examine whether Reelin affects cilia length in interneurons, we measured cilia length in WT and *reeler* mice of P30 using AC3 immunostaining in combination with interneuron-specific markers GAD65 (Kajita and Mushiake, 2021). In the CA1 region of the hippocampus, there was no significant difference in cilia length of interneurons between genotypes (Fig. S6F-G). Similarly, in both the deep and superficial layers of the neocortex, cilia length in interneurons did not differ between WT and *reeler* (Table S4). These findings demonstrate that Reelin does not influence primary cilia length in interneurons.

Lastly, we measured cilia length of astrocytes in WT and *reeler* mice of P30, using Arl13b antibody (Sterpka and Chen, 2018) co-immunostaining with a classical marker GFAP antibody. In the CA1 region of the hippocampus, there was no significant difference in cilia length between genotypes (Fig. S6 H-I). Similarly, in both the deep and superficial layers of the neocortex, GFAP-positive astrocytic cilia length did not differ between WT and *reeler* mice (Table S4). These findings indicate that Reelin does not affect cilia length in GFAP-positive astrocytes.

## Discussion

### The Apical Localization and Directional Orientation of Primary Cilia may be Linked to Neuronal Polarity

This study identifies alterations in the directional orientation, apical localization, and length of primary cilia in cortical principal neurons of *reeler* mice. Along with previous work reporting reverse soma movement of pyramidal neurons (Yang et al., 2025), these results suggest that Reelin-directed neuronal placement is associated with the establishment of the apical localization and directional orientation of cilia in principal neurons. This study also highlights primary cilia as an apical domain of pyramidal neurons in the cerebral cortex.

It seems evident how Reelin affects the subcellular localization of cilia in principal neurons. Reelin signaling plays a critical role in determining the subcellular localization of the centrosome in principal neurons in the cerebral cortex. As neurons complete their radial migration, Reelin, through activation of its receptors and subsequent downstream signaling, directs soma translocation and the positioning of the centrosome at the base of the apical dendrite (the apical pole) (Chai and Frotscher, 2016; Cooper, 2008; Jossin, 2020; Trommsdorff et al., 1999). Because the primary cilium is derived from the mother centriole, the positioning of the centrosome in the apical pole provides the structural basis for the localization of the cilium to the apical side of the neuronal soma.

The functional significance of positioning pia-oriented cilia on the apical side of pyramidal neurons is yet to be explored. Primary cilia function as specialized signaling compartments containing numerous receptors, including G protein-coupled receptors (Mukhopadhyay and Rohatgi, 2014; Pal and Mukhopadhyay, 2015; Tschaikner et al., 2020; Wang and Ogden, 2025). The preferential localization of the cilium at the base of the apical dendrite spatially distant this signaling domain from AIS, the site of action potential initiation for electric output. Interestingly, the cilia-at-apical and AIS-at-basal pattern is the strongest in late-born pyramidal neurons (Fig. 1 and S1). Such spatial pattern does not exist in atypical pyramidal neurons, excitatory neurons of non-laminated brain regions, or interneurons. The separation of the ciliary domain from the axonal output may therefore represent an additional feature of pyramidal neuron polarity, which may enhance direction information flow from the apical dendrite to basal output (Larkum, 2013; Spruston, 2008), at least during early postnatal neurodevelopment. Moreover, pia-oriented primary cilia at the apical side are closer to apical synapses than to synapses on basal dendrites, preferentially facilitating signal integration from apical synapses (Ott et al., 2024; Sheu et al., 2022; Wu et al., 2024). However, whether this spatial pattern has functional consequences to enhance neuronal polarity remains to be experimentally determined.

Recent observations using electron microscopy (EM) in the human temporal cortex, excised out of a 45-year-old seizure patient (Wu et al., 2024), and in the mouse visual cortex (Ott et al., 2024) provide additional evidence that the primary cilia of principal neurons are spatially defined rather than randomly positioned. Wu et al. reported that most neuronal primary cilia in the human temporal lobe are oriented approximately parallel to the cortical wall and preferentially localize toward dendritic rather than axonal domains (Ott et al., 2024). However, this study did not identify the preferential pia-directed orientation observed in our studies. Possible explanations for this difference include the age and epileptic state of the patient. Our analyses were performed on mouse brain samples of P5-P30, during which postnatal neuronal repositioning occurs. Around P20, the primary cilia of pyramidal neurons exhibit the strongest pia-oriented preference (Yang et al., 2025). This pia-orientation preference gradually becomes less pronounced as age (Yang et al., 2025) and can change under epileptic state (Kirschen et al., 2017).

### How does Reelin Influence Cilia Directional Orientation?

This report, along with our prior study (Yang et al., 2025), supports a working model in which pia-directed cilia orientation is not automatically formed when cilia are expressed, but gradually emerged from the postnatal reverse movement. During embryonic neurodevelopment, Reelin coordinates centrosome positioning within the leading process, which later become the apical dendrite, to lead rapid radial migration (Jossin, 2020). Neurons are directed to migrate along radial fibers outward the pia and then stop upon encountering Reelin (Chai and Frotscher, 2016; Cooper, 2008; Frotscher et al., 2009; Rice and Curran, 2001), resulting in their accumulation in the upper layers beneath the marginal zone (Fig. 10). Following birth, the high cell density or crowdedness in the upper layers may drive the slow reverse soma movement of late-born neurons toward the deeper regions age (Yang et al., 2025). As the somata move backwards, the centrioles and expressed cilia follow, resulting in the cilia becoming oriented towards the pia. Early-born neurons may be indirectly influenced by the movement of late-born neurons. In the r*eeler* cerebral cortex, where embryonic radial migration is severely disrupted and late-born neurons do not migrate toward or accumulate in the superficial layer. This may affect postnatal reverse movement and secondly cilia orientation (Fig. 10). However, the partial postnatal correction observed in *reeler* late-born neocortical neurons suggests that Reelin is unlikely to be the sole determinant of cilia directionality. Hypothetically, some intrinsic developmental programs in late-born pyramidal neurons, interactions with neighboring cells or extracellular matrix may also contribute to the orientation of primary cilia.

**Fig. 10.**
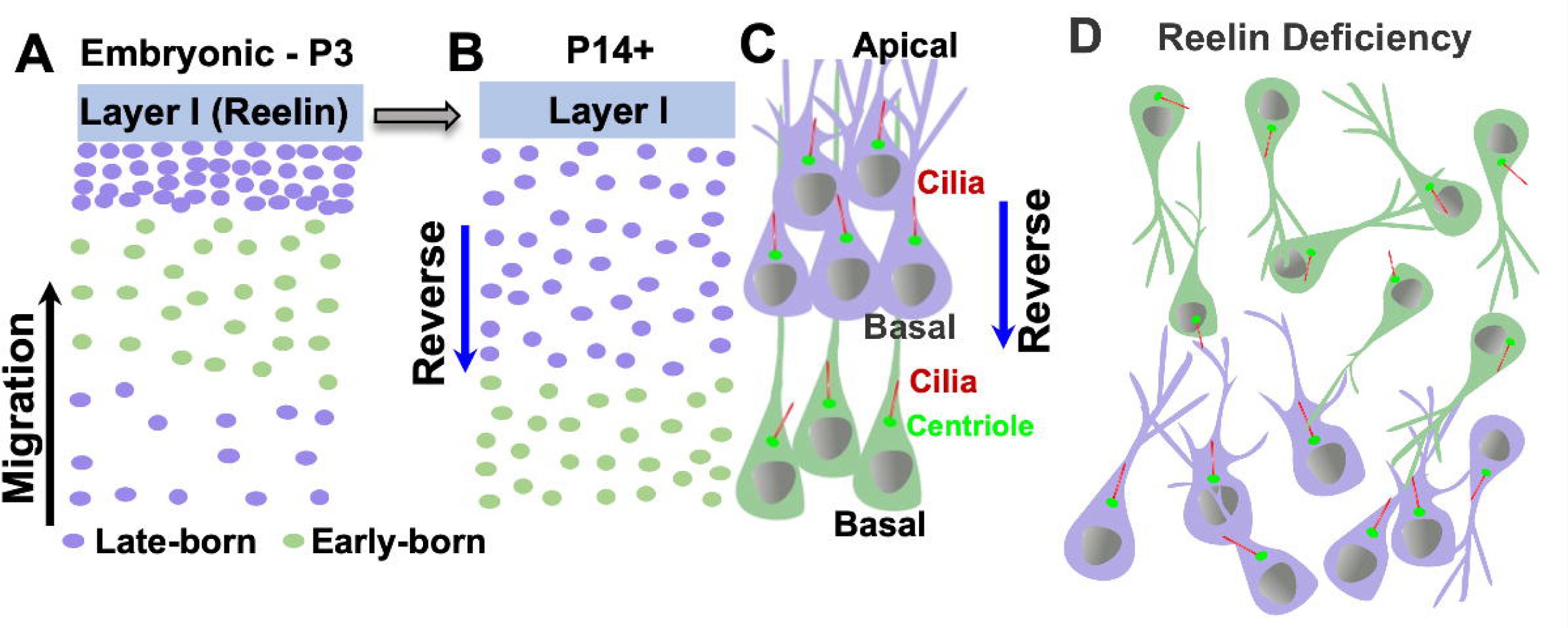
Working model of how Reelin affects directional orientation of cilia in the neocortex. (**A**) During embryonic migration, the centrosomes lead the radial migration without cilia expression. Following the completion of radial migration, late-born neurons accumulate densely in the upper layer immediately beneath the marginal zone (Layer I). (**B**) A slow reverse movement is required for postnatal positioning and sparsely redistribute neurons to the deeper layers from P3 until P14, cilia are expressed during this period. **(C)** As neuronal somata move backward, the centrioles and emerged cilia follow, resulting in the cilia becoming oriented towards the pia. (**D**) In the absence of Reelin, neocortical neurons fail to establish the normal inside-out lamination pattern and are unable to densely accumulate in the upper layer. Consequently, this impairs reverse movement and leaves neurons unable to maintain correct re-positioning direction, causing them to drift aimlessly. This aberrant movement ultimately alters the directional orientation of cilia. A portion of late-born neurons however still reverse to some degree, thereby manifesting postnatal correction in cilia orientation.

### Distinct Cilia Phenotypes among Neuronal Populations in *reeler*

Reelin deficiency does not affect cilia pattern uniformly across cerebral cortical regions. In the DG, CA3, and piriform cortex, where laminated structure is severely disrupted in *reeler*, primary cilia of principal neurons show marked loss of apical localization and directional orientation. A similarly severe phenotype is observed among early-born pyramidal neurons in the neocortex. In contrast, late-born neurons in the neocortex and hippocampal CA1 exbibit partial restoration during postnatal development.

One question arises why late-born pyramidal neurons in inside-out laminated region retain a greater capacity for cilia apical location and pia-orientation in the absence of Reelin. One possibility is that late-born neurons emerge later in cortical evolution and development and may acquire programs that confer greater resilience to developmental alterations. Late-born neurons may evolve to develop an intrinsic propensity for postnatal reverse movement and consequently re-alignment of cilia orientation, even in the absence of Reelin. In contrast, principal neurons in evolutionarily conserved cortical regions do not reverse, potentially accounting for the less pronounced postnatal correction. Thus, rather than being established exclusively during embryonic migration, the polarity of late-born neurons may continue to be refined during postnatal development. In this context, postnatal reverse soma movement could contribute not only to the refinement of neuronal position and cortical layering (Yang et al., 2025), but also to the enhancement of neuronal polarity and direction information flow. However, experimental testing is needed to evaluate this possibility.

### Reelin Deficiency Increases Cilia Length in Principal Neurons in the Cerebral Cortex

In addition to changes in cilia localization and orientation, Reelin deficiency also leads to increased cilia length. Primary cilia of principal neurons in the mouse cerebral cortex emerge predominantly between P3 and P14, after which their length remains relatively stable in WT mice (Yang et al., 2025). In *reeler*, however, primary cilia become significantly longer across multiple cortical regions, with this phenotype becoming apparent after P14 (Table S4).

Reelin has a particularly important role in neuronal placement within cerebral cortical regions, where principal neurons typically exhibit directional cilia orientation, and loss of Reelin increase cilia length. In contrast, neurons in nucleated brain regions generally lack cilia directionality or preference for apical cilia localization age (Yang et al., 2025), and Reelin deficiency has no effect on cilia length. Consistently, cilia length in *Reln*+/− mice, which retain largely normal cortical lamination (Joly-Amado et al., 2023; Tueting et al., 1999), is not markedly different from that of WT mice. These observations suggest that cilia elongation is associated with severe disruption of Reelin signaling and cortical lamination rather than with reduced Reelin dosage.

The biological significance of this elongation remains unknown. Primary cilia are specialized signaling compartments, and changes in cilium length potentially alters the surface area, ciliary volume and its interactions with the extracellular environment. Appropriate regulation of cilium length may therefore influence the strength of ciliary signaling (Macarelli et al., 2023; Wang and Ogden, 2025). Nevertheless, we do not know how Reelin signaling regulates cilia length. Cilia elongation in *reeler* could arise secondarily from altered neuronal migration. Distinguishing these possibilities will require temporal and cell-specific manipulation of Reelin signaling and *in vitro* cell culture study.

### Limitations of this Study

Several limitations should be considered when interpreting these findings. First, although there is a strong association between Reelin deficiency and alterations in ciliation pattern, the use of a global Reelin-null mutant cannot distinguish direct effects of Reelin signaling on primary cilia from secondary consequences of abnormal neuronal migration. Second, this study is primarily morphological and does not establish a direct causal relationship between altered cilia pattern and Reelin signaling. Future studies using temporally controlled, cell-type-specific manipulation of Reelin signaling are needed to distinguish its embryonic effects neuronal migration from secondary effects on postnatal re-positioning, refinement of neuronal polarity. *Ex vivo* time-lapse cilia-labeled imaging on organotypic slice culture will assess the causal relationship between reverse movement and cilia orientation. These experiments may also help test the hypothesis that Reelin signaling establishes the general framework for cortical lamination during embryonic development, while postnatal repositioning further promotes the neuronal polarity in pyramidal neurons, in part through apically localized, pia-oriented primary cilia.

## Funding and Acknowledgments

This research was supported by National Institutes of Health Grants R15MH126317 and R15MH125305 to X.C.; and by UNH CoRE awards, Cole Neuroscience and Behavioral Faculty Research Awards, and a COLSA Community of Teaching and Research Scholar Award (to X.C.); Grants-in-aid for Scientific Research of Japan Society for Promotion of Science JP24KJ1889 to Y.T.; Grants-in-aid for Scientific Research of Japan Society for Promotion of Science JP20H03384 and JP23K27324 (to M.H).; Grant-in-Aid for Outstanding Research Group Support Program in Nagoya City University (Grant Number 2401101). S.A. and S.M. were funded by UNH Summer TA Research Fellowships (STAF), K.B. by a McNair Scholar Award, and K.J. by an Undergraduate Research Award at UNH. We are grateful to the University Instrumentation Center at UNH for confocal imaging service.

## Supplemental Figure Legends

**Fig. S1. Cilia-at-apical and AIS-at-basal prominent pattern in the upper layer of the neocortex**. Representative images showing pia-oriented cilia and apical localization in the neocortex of P30. AIS and cilia were marked by Ankyrin G and AC3 respectively. The pattern of cilia-at-apical and AIS-at-basal (with examples encircled in white) is stronger in the upper layers (**Top**) than in the deep layer (**Bottom**).

**Fig. S2. Lack of directional cilia orientation and apical cilia location in atypical pyramidal neurons in the WT deep neocortex. (A)** Representative cilia images of atypical pyramidal neurons stained with NeuN, AC3 and Pericentrin antibodies. (**B**) Glutaminase I staining was used to confirm excitatory neurons. **(C)** Representative images of cilia and AIS in atypical pyramidal neurons (marked by Ctip2) in the deep cortical layers. The examples show cilia and AIS (indicated by white arrows) in reference to the nuclei, which are randomly positioned and lack the distal-opposite alignment characteristic of late-born cortical neurons. **(D)** Histogram and polar plot showing the angle frequency distribution of cilia orientation (0° - 360°), with a flattened fitting curve indicating lack of directionality. **(E)** Cilia location in 4 quadrants of atypical pyramidal neurons. One-way ANOVA test.

**Fig. S3. (A-D) Primary cilia of excitatory neurons in the WT amygdala lack directional orientation or preference for subcellular localization. (A)** Representative images of cilia (marked by AC3) and AIS (marked by Ankyrin G) in the amygdala show irregular cilia orientation and random cilia localization in the soma, which lack the typical alignment distal to the AIS. **(B)** Representative images of excitatory neurons, marked by a Glutaminase antibody, in the WT amygdala. **(C)** Cilia orientation histogram and polar plot show the distribution of cilia orientation angles of excitatory neurons in the WT amygdala. **(D)** cilia location in four quadrants of amygdala excitatory neurons. One-way ANOVA test. **(E-F)** Primary cilia lack preference to location in interneurons and GFAP-positive astrocytes. (Left Insert) Diagrams showing how 4 quadrants in the soma (Top, Bottom, Left and Right) were defined in reference to the nucleus. Interneurons and astrocytes were marked by GAD65 and GFAP antibodies, respectively. One-way ANOVA test.

**Fig. S4. *reeler* Late-born neurons maintain a stronger tendency than early-born neurons to re-align cilia to the apical side.** Relative position of neuronal primary cilia (marked by AC3), soma (marked by Sabt2 or Ctip2), and AIS (marked by Ankyrin G) in late-born (**A**) and early-born neurons (**B**) of WT and *reeler* neocortex at P10 and P30.

**Fig. S5. Reelin haploinsufficiency does not affect cilia length in principal neurons in the cerebral cortex.** Representative cilia images and cilia length in the CA1 (**A**), superficial neocortex (**B**), and DG (**C**) of WT and Reelin +/- mice revealed no differences between WT and Reelin +/- mice. Un-paired Welch’s T-tests, ns, not significant.

**Fig. S6. (A-E) Reelin does not influence cilia length in non-laminated brain regions.** (**A**) Allen brain atlas showing brain region examined. **(B)** Representative neuronal cilia images and scatter plot showing comparable cilia length in the thalamus of WT and *reeler* mice of P30. Similar measurements in the striatum **(C)**, hypothalamus **(D)**, and amygdala **(E)** also showed no significant differences between genotypes. Two-tailed unpaired Welch’s T-tests, ns, not significant. **(F-I) Reelin does not affect cilia length in interneurons or GFAP-positive astrocytes. (F)** Representative images of individual interneurons in the WT and *reeler* neocortex of P30. **(G)** Scatter plots of primary cilia length in interneurons of WT and *reeler* mice revealed no significant differences between genotypes. Interneurons were identified by GAD65 antibody. **(H)** Representative confocal images of individual GFAP-positive astrocytes in the WT and Reeler hippocampal CA1 region of P30. **(I)** Scatter graphs of primary cilia length in GFAP-positive astrocytes of WT and *reeler* mice revealed no significant differences between genotypes. Statistical analyses were performed using Welch’s T-test. At P30, Arl13b antibody generally labels GFAP-positive astrocytic primary cilia, not neuronal cilia. A single cilium was assigned to an astrocyte based on its proximity to the nucleus and GFAP staining. Table S4 summarizes cilia length of various cell types in the different regions. Two-tailed unpaired Welch’s T-tests, ns, not significant.

---

**Table S1.** Primary and secondary antibodies used for immunofluorescence staining.

| Antibody/target | Host species | Antibody type | Manufacturer | Catalog number | Working dilution |
| --- | --- | --- | --- | --- | --- |
| Anti-AC3 | Rabbit | Primary | EnCor Biotechnology Inc. | RPCA-AC3 | 1:5,000 |
| Anti-AC3 | Chicken | Primary | EnCor Biotechnology Inc. | CPCA-ACIII | 1:5,000 |
| Anti-Pericentrin | Rabbit | Primary | Abcam | ab4448 | 1:500 |
| Anti-Glutaminase 1 (GLS1) | Rabbit | Primary | Synaptic Systems | 456 003 | 1:500 |
| Anti-Ctip2 | Rat | Primary | Abcam | ab18465 | 1:500 |
| Anti-Satb2 | Rabbit | Primary | Abcam | ab92446 | 1:500 |
| Anti-Ankyrin G | Mouse | Primary | Synaptic Systems | 181 011 | 1:500 |
| Anti-NeuN, clone A60 | Mouse | Primary | Millipore Sigma | MAB377 | 1:500 |
| Anti-GAD65 | Rabbit | Primary | Proteintech | 21760-1-AP | 1:500 |
| Anti-GFAP | Rabbit | Primary | Dako | Z0334 | 1:500 |
| Donkey anti-rabbit IgG, Alexa Fluor 546 | Donkey | Secondary | Invitrogen | A10040 | 1:1000 |
| Donkey anti-rabbit IgG, Alexa Fluor 647 | Donkey | Secondary | Invitrogen | A31573 | 1:1000 |
| Donkey anti-chicken IgY, Alexa Fluor 568 | Donkey | Secondary | Invitrogen | A78950 | 1:1000 |
| Donkey anti-rat IgG, Alexa Fluor 488 | Donkey | Secondary | Invitrogen | A21208 | 1:1000 |
| Donkey anti-rat IgG, Alexa Fluor 647 | Donkey | Secondary | Jackson Immuno Research | 712-605-150 | 1:1000 |
| Donkey anti-mouse IgG, Alexa Fluor 488 | Donkey | Secondary | Invitrogen | A21202 | 1:1000 |

**Table S2.**
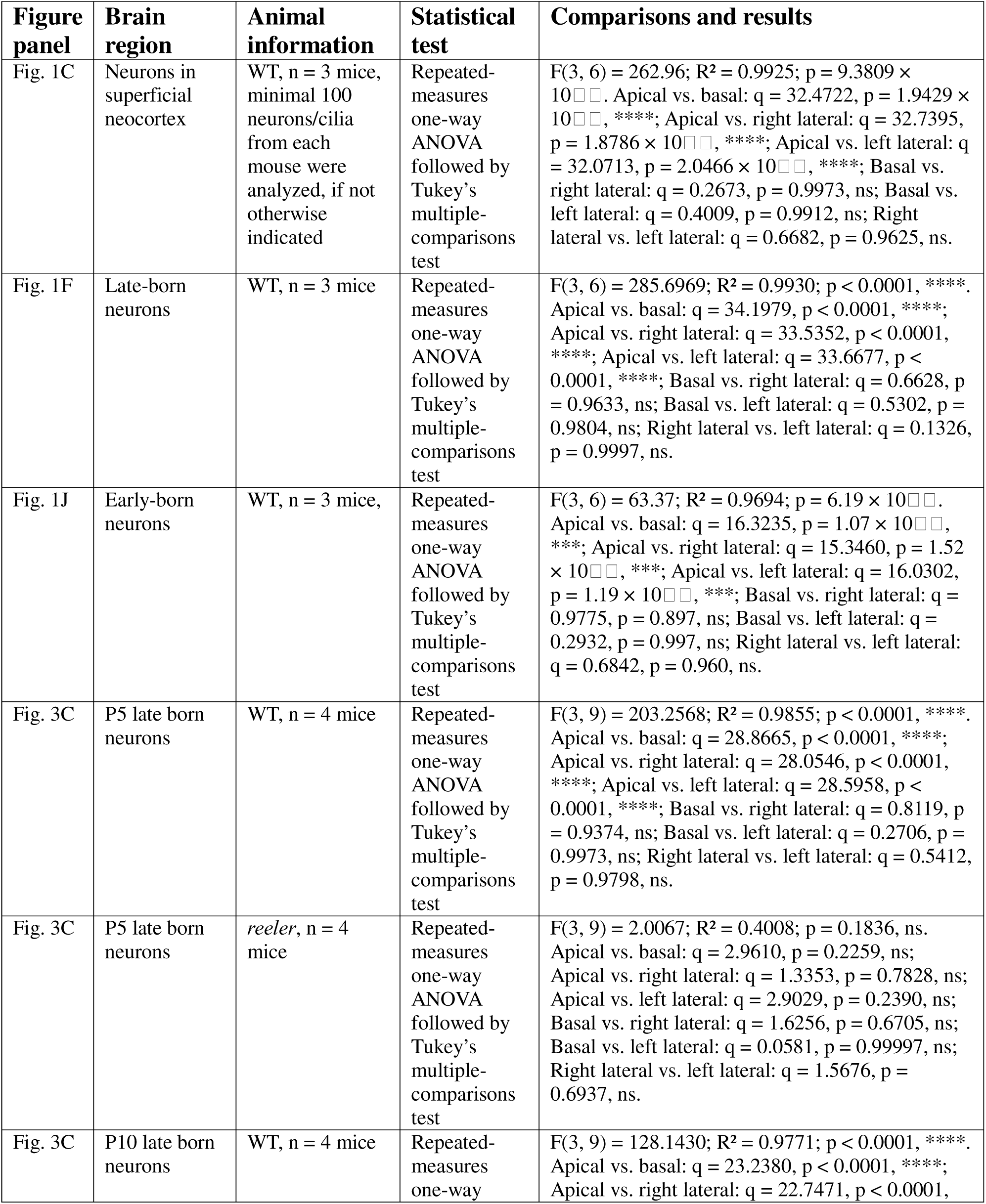
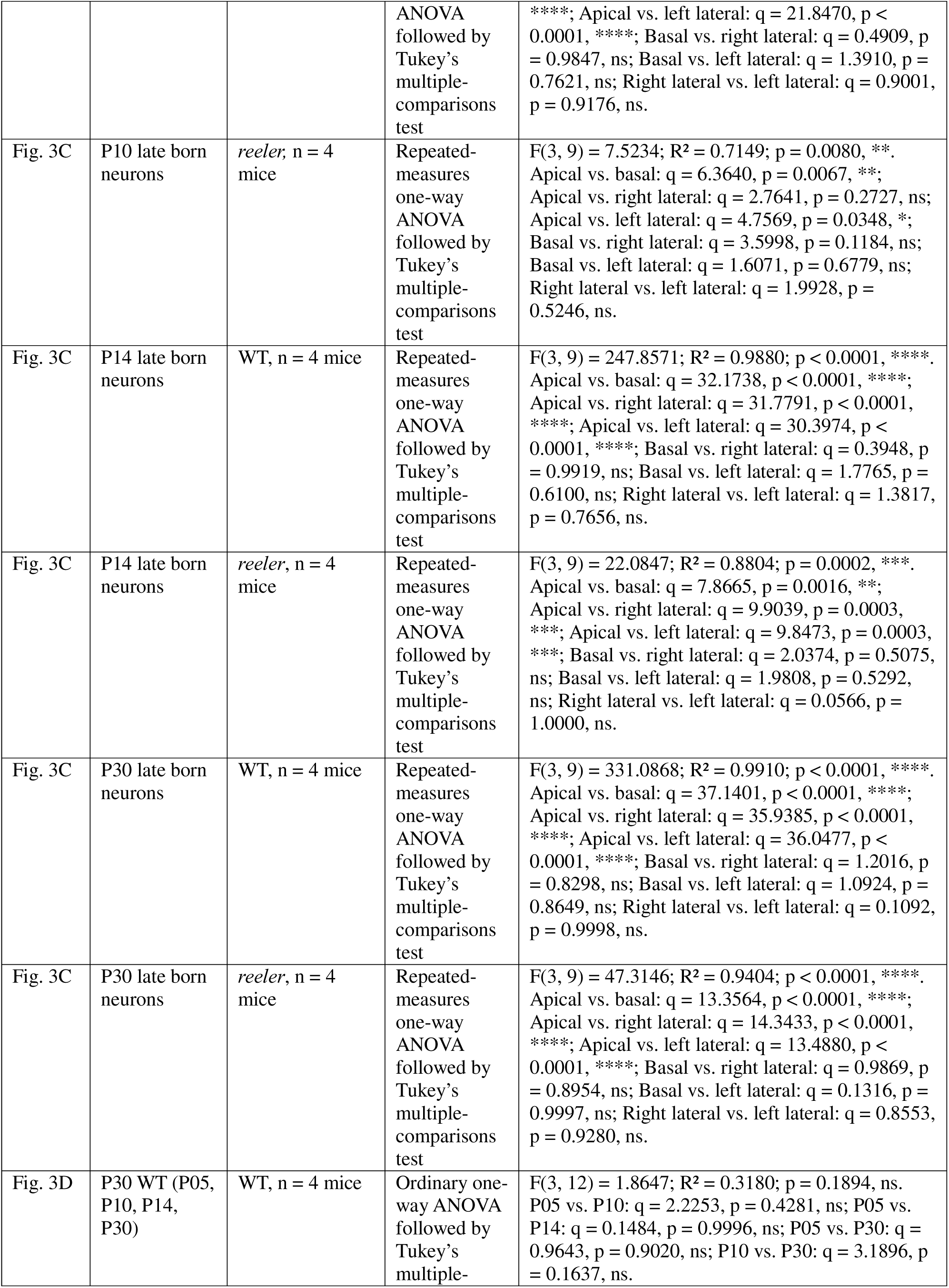
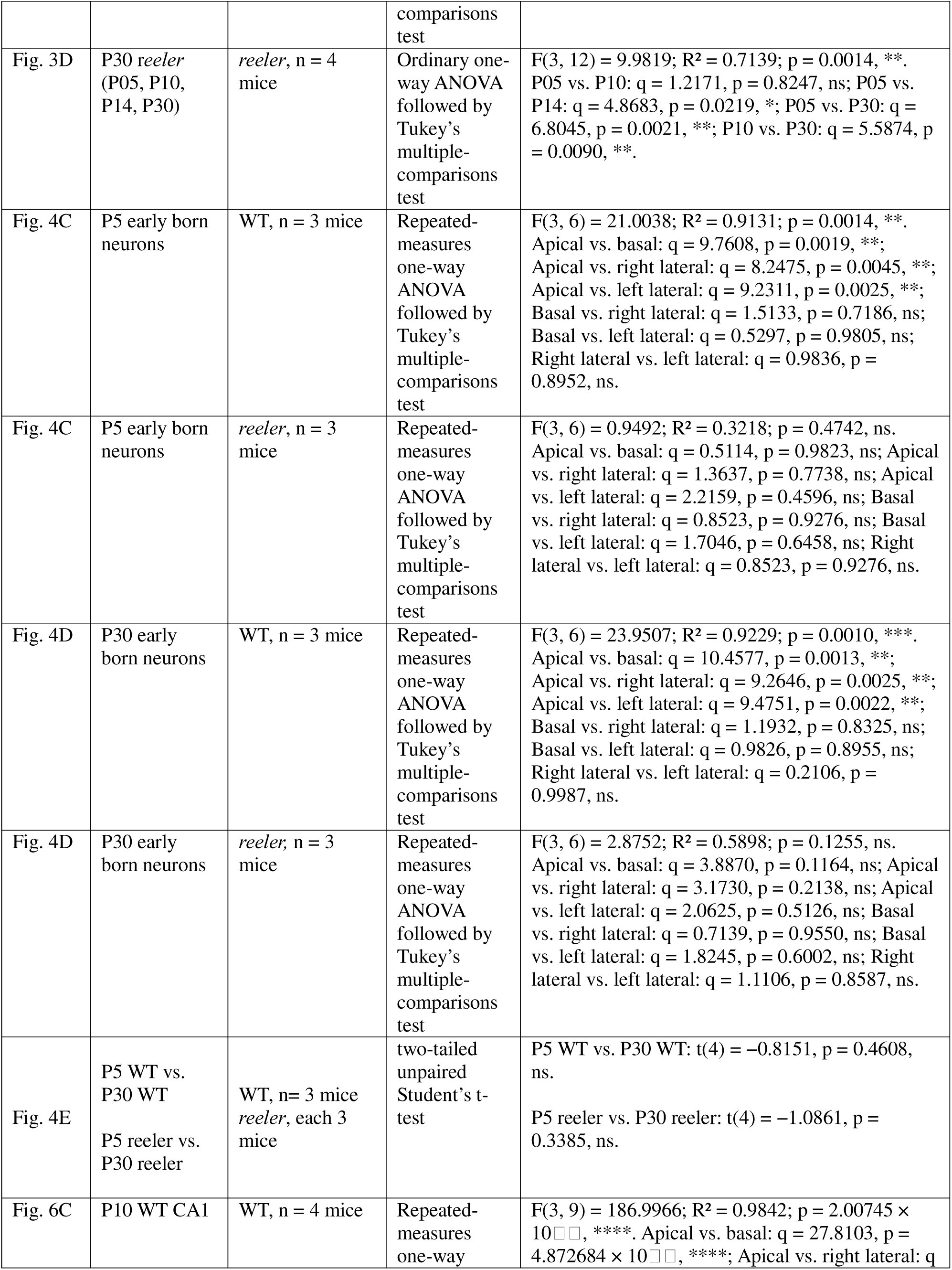
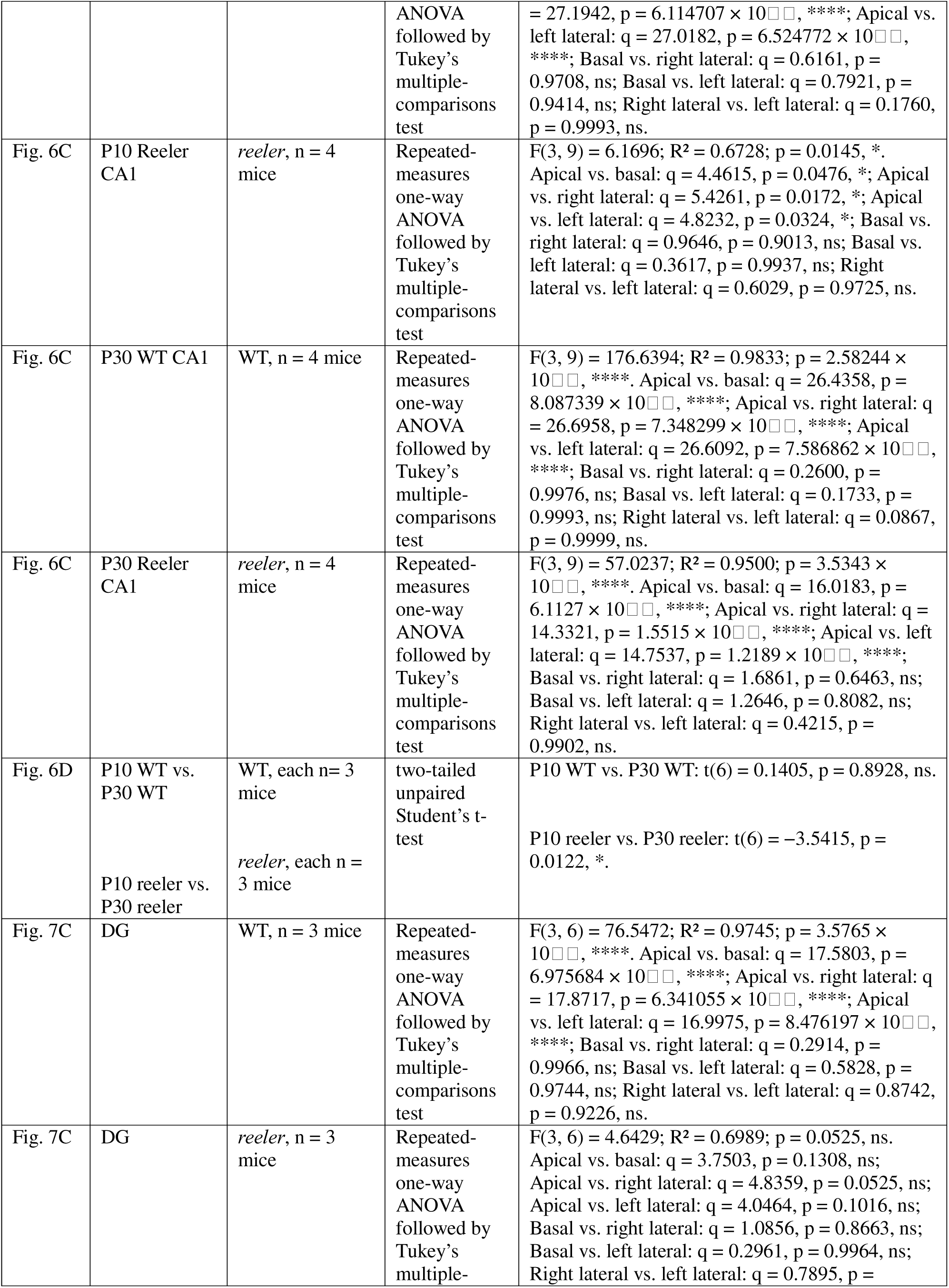
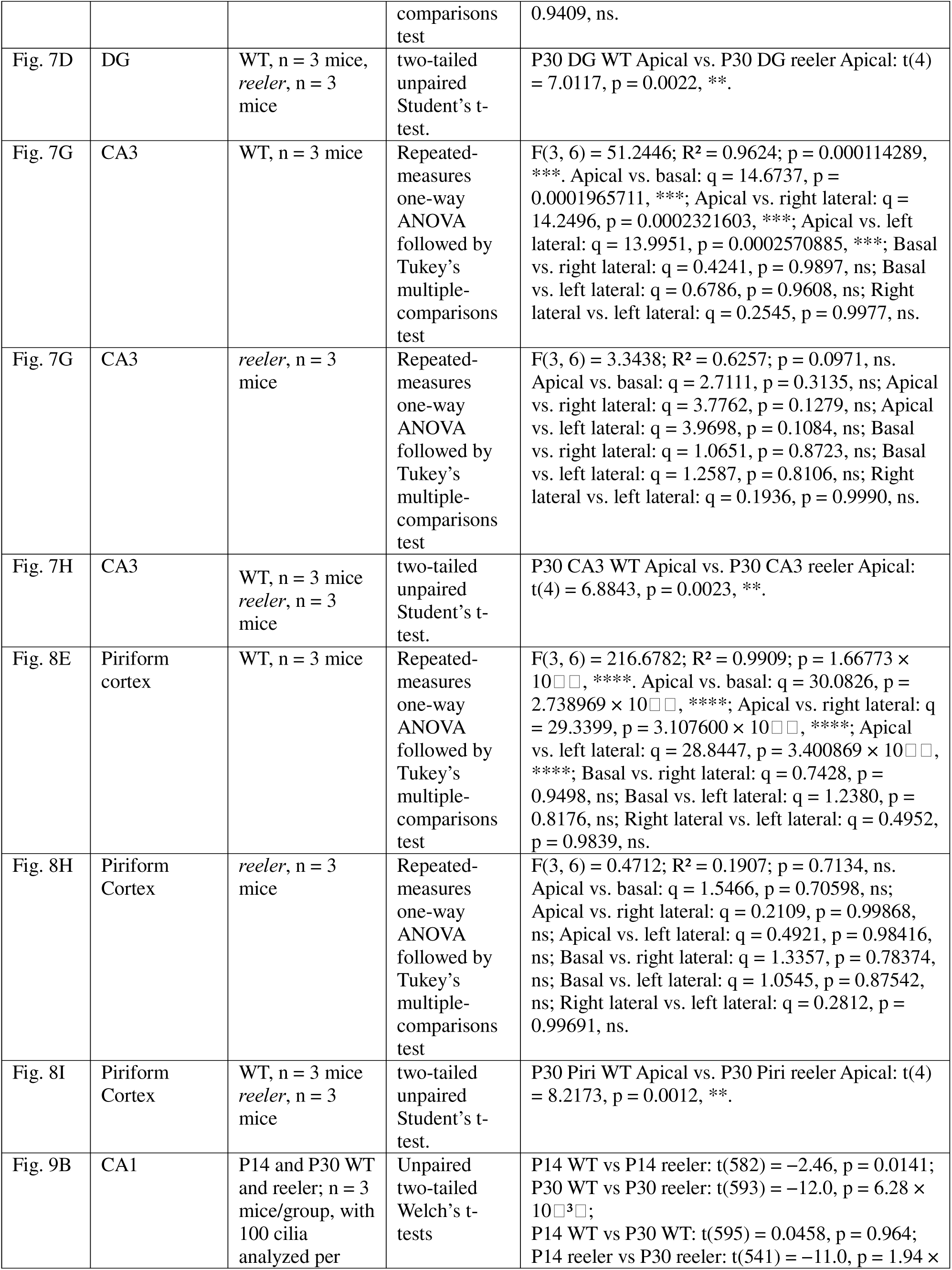
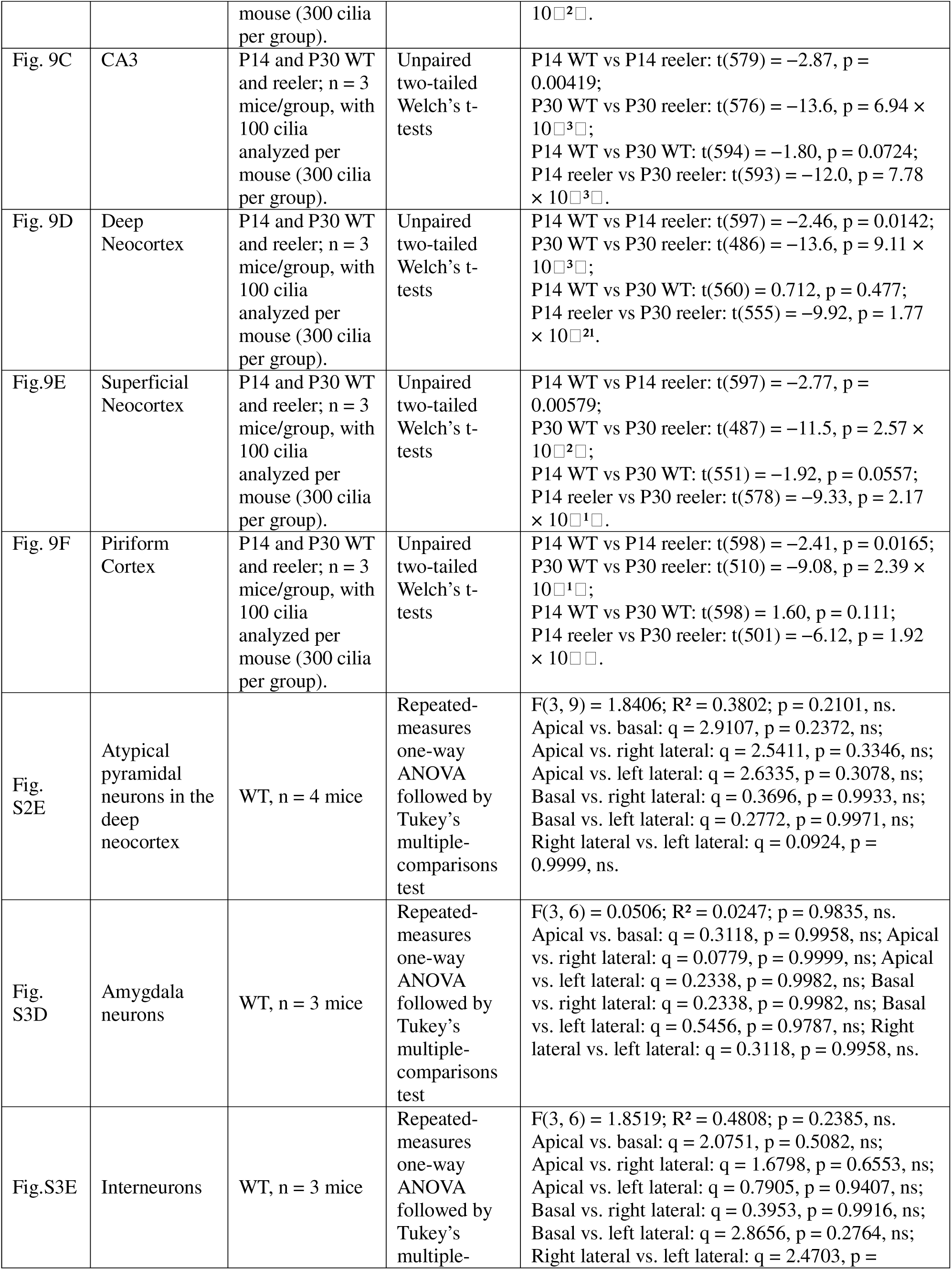
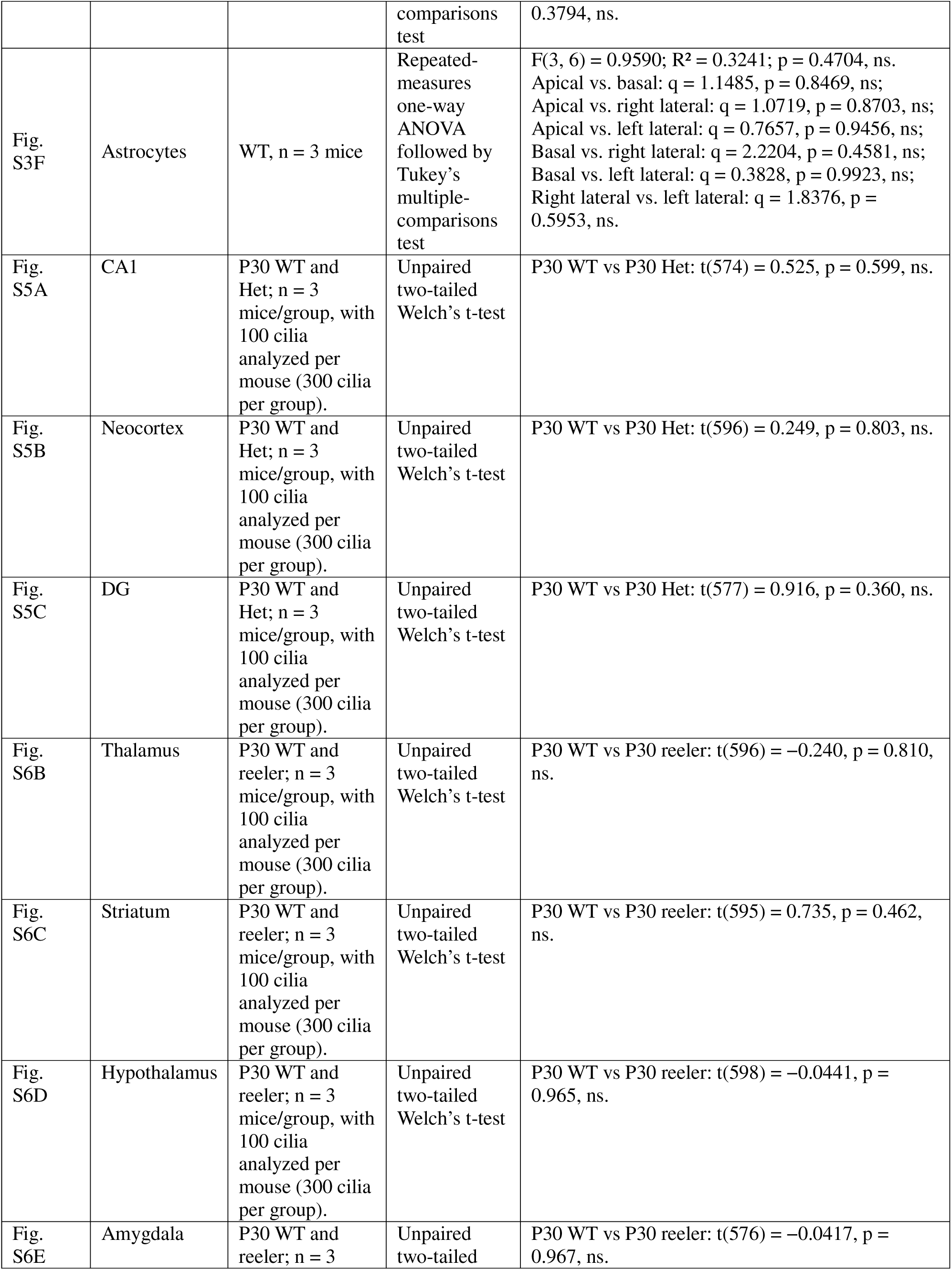
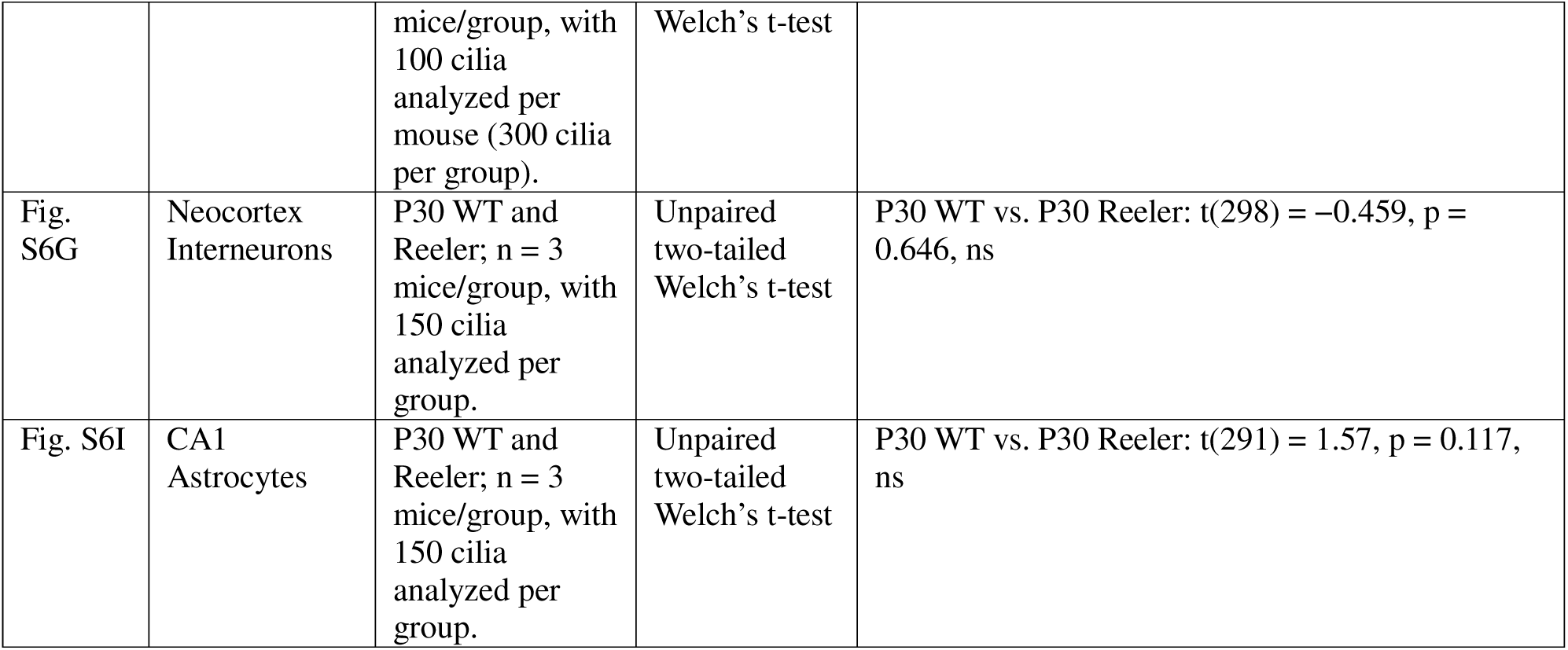
Detailed Information of Statistical Tests.

**Table S3:** Cilia Orientation and Subcellular Location in Brain Regions of WT and *reeler* Mice.

| Brain Regions<br>or Cell Types | WT (P30) |  |  | *reeler* (P30) |  |  |
| --- | --- | --- | --- | --- | --- | --- |
|  | Cilia<br>Orientation | Lemina<br>Cell Density | Cilia Location | Cilia<br>Orientation | Lemina<br>Cell Density | Cilia Location |
| DG | Roughly<br>opposite | Compact | Mainly apical<br>quadrant | Lack specific<br>direction | Disorganized | Broadly<br>distributed |
| CA3 | Weakly<br>opposite | Compact | Mainly apical<br>quadrant | Lack specific<br>direction | Disorganized | Broadly<br>distributed |
| Dorsal CA1 | mainly toward<br>SR | Compact | Apical quadrant | Opposite | Compact, Split<br>SP | Mainly apical<br>quadrant |
| Ventral CA1 | mainly toward<br>SR | Compact | Apical quadrant | Opposite | Compact, Split<br>SP | Mainly apical<br>quadrant |
| Neocortex<br>(Late-born) | Oriented to the<br>pia | Sparse | Apical quadrant | Mainly<br>oriented to the<br>pia | Inverted,<br>partially affected | Mainly apical<br>quadrant |
| Neocortex<br>(Early-born) | Oriented to the<br>pia | Sparse | Mainly apical<br>quadrant | Lack specific<br>direction | Inverted,<br>Disorganized | Broadly<br>distributed |
| Piriform Cortex<br>(Layer II) | Opposite | Compact | Mostly apical<br>quadrant | Lack specific<br>direction | Disorganized | Broadly<br>distributed |
| Piriform Cortex<br>(Layer III) | Oriented to the<br>pia | Sparse | Apical quadrant | Lack specific<br>direction | Disorganized | Broadly<br>distributed |
| CA1<br>Interneuron | Lack specific<br>direction | Not layered | Broadly<br>distributed | Lack specific<br>direction | Not layered | Broadly<br>distributed |
| Neocortex<br>Interneuron | Lack specific<br>direction | Not layered | Broadly<br>distributed | Lack specific<br>direction | Not layered | Broadly<br>distributed |
| CA1 GFAP-positive<br>Astrocyte | Lack specific<br>direction | Not layered | Broadly<br>distributed | Lack specific<br>direction | Not layered | Broadly<br>distributed |
| Neocortex,<br>GFAP-positive<br>Astrocyte | Lack specific<br>direction | Not layered | Broadly<br>distributed | Lack specific<br>direction | Not layered | Broadly<br>distributed |
| Thalamus | Lack specific<br>direction | Not layered | Broadly<br>distributed | Lack specific<br>direction | Not layered | Broadly<br>distributed |
| Hypothalamus | Lack specific<br>direction | Not layered | Broadly<br>distributed | Lack specific<br>direction | Not layered | Broadly<br>distributed |
| Amygdala | Lack specific<br>direction | Not layered | Broadly<br>distributed | Lack specific<br>direction | Not layered | Broadly<br>distributed |
| Striatum | Lack specific<br>direction | Not layered | Broadly<br>distributed | Lack specific<br>direction | Not layered | Broadly<br>distributed |

**Table S4:** Cilia Length in Various Brain Regions and Cell Types of WT and *reeler* Mice.

| Brain Region | Layering | P14 |  | P30 |  |
| --- | --- | --- | --- | --- | --- |
|  |  | WT (μm) | *reeler* (μm) | WT (μm) | *reeler* (μm) |
| CA1 | Laminated | 5.21 ± 0.21 | 5.86 ± 0.10 (*) | 5.45 ± 0.23 | 9.45 ± 0.46 (**) |
| CA3 | Laminated | 5.13 ± 0.14 | 5.72 ± 0.08 (*) | 5.32 ± 0.15 | 6.71 ± 0.29 (*) |
| DG | Laminated | 2.82 ± 0.13 | 3.64 ± 0.16 (**) | 2.90 ± 0.15 | 5.02 ± 0.20 (***) |
| Deep Neocortex | Laminated | 4.31 ± 0.10 | 4.82 ± 0.20 (*) | 4.49 ± 0.17 | 5.92 ± 0.10 (***) |
| Superficial Neocortex | Laminated | 5.23 ± 0.11 | 5.71 ± 0.07 (**) | 5.36 ± 0.07 | 6.88 ± 0.20 (***) |
| Piriform Cortex | Laminated | 4.61 ± 0.25 | 5.12 ± 0.13 (*) | 4.83 ± 0.09 | 6.02 ± 0.27 (**) |
| Entorhinal Cortex | Laminated | 4.80 ± 0.27 | 5.33 ± 0.11 (*) | 5.15 ± 0.14 | 6.01 ± 0.08 (***) |
| Thalamus | Non-laminated | ND | ND | 5.53 ± 0.32 | 5.53 ± 0.34 (n.s.) |
| Hypothalamus | Non-laminated | ND | ND | 5.48 ± 0.35 | 5.53 ± 0.33 (n.s.) |
| Amygdala | Non-laminated | ND | ND | 5.28 ± 0.14 | 5.29 ± 0.16 (n.s.) |
| Striatum | Non-laminated | ND | ND | 6.41 ± 1.02 | 6.43 ± 1.01 (n.s.) |
| GFAP-positive Astrocyte in CA1 | Non-laminated | ND | ND | 5.12 ± 0.01 | 5.14 ± 0.03 (n.s.) |
| GFAP-positive Astrocyte in deep cortex | Non-laminated | ND | ND | 4.25 ± 0.14 | 4.34 ± 0.11 (n.s.) |
| GFAP-positive Astrocyte in superficial cortex | Non-laminated | ND | ND | 4.51 ± 0.07 | 4.50 ± 0.06 (n.s.) |
| Interneuron CA1 | Non-laminated | ND | ND | 5.09 ± 0.11 | 5.13 ± 0.14 (n.s.) |
| Interneuron Deep Cortex | Non-laminated | ND | ND | 5.13 ± 0.19 | 5.11 ± 0.10 (n.s.) |
| Interneuron Superficial Cortex | Non-laminated | ND | ND | 5.16 ± 0.34 | 5.17 ± 0.31 (n.s.) |

