## Supplementary figures and images for "Reelin Deficiency Alters the Directional Orientation, Apical Localization and Length of Primary Cilia in Principal Neurons in the Cerebral Cortex"

### Supplemental Figure 1-6

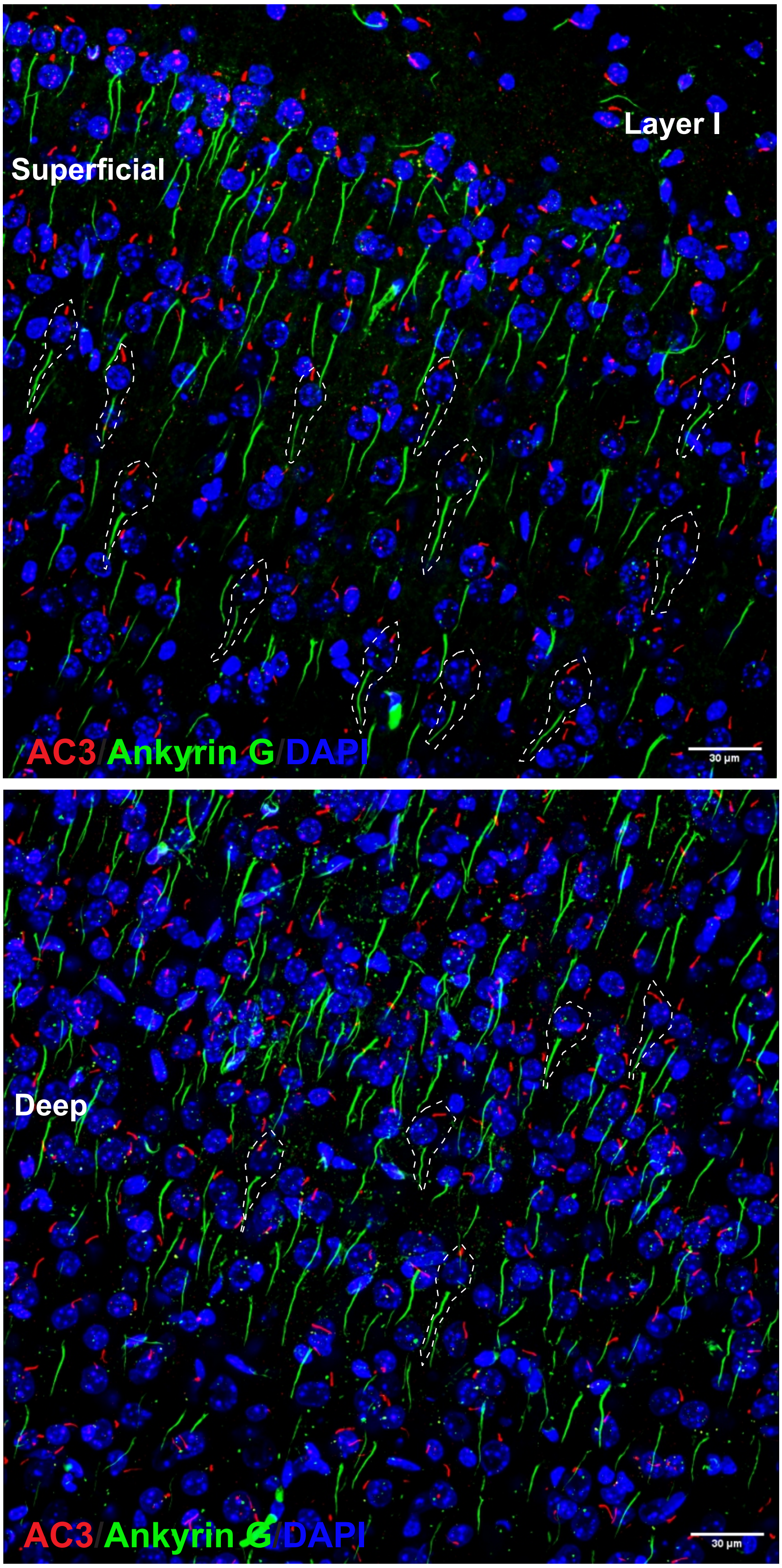

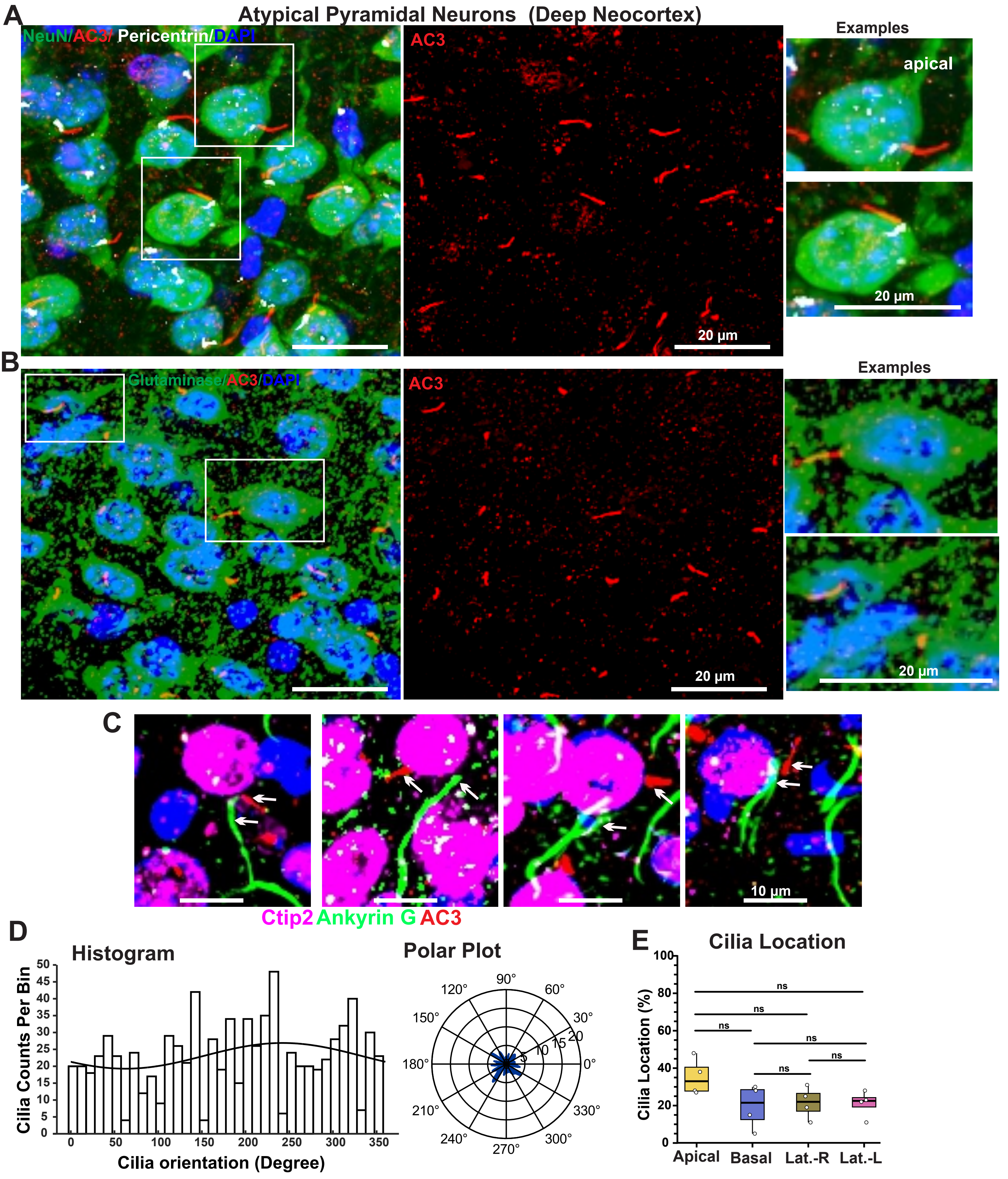

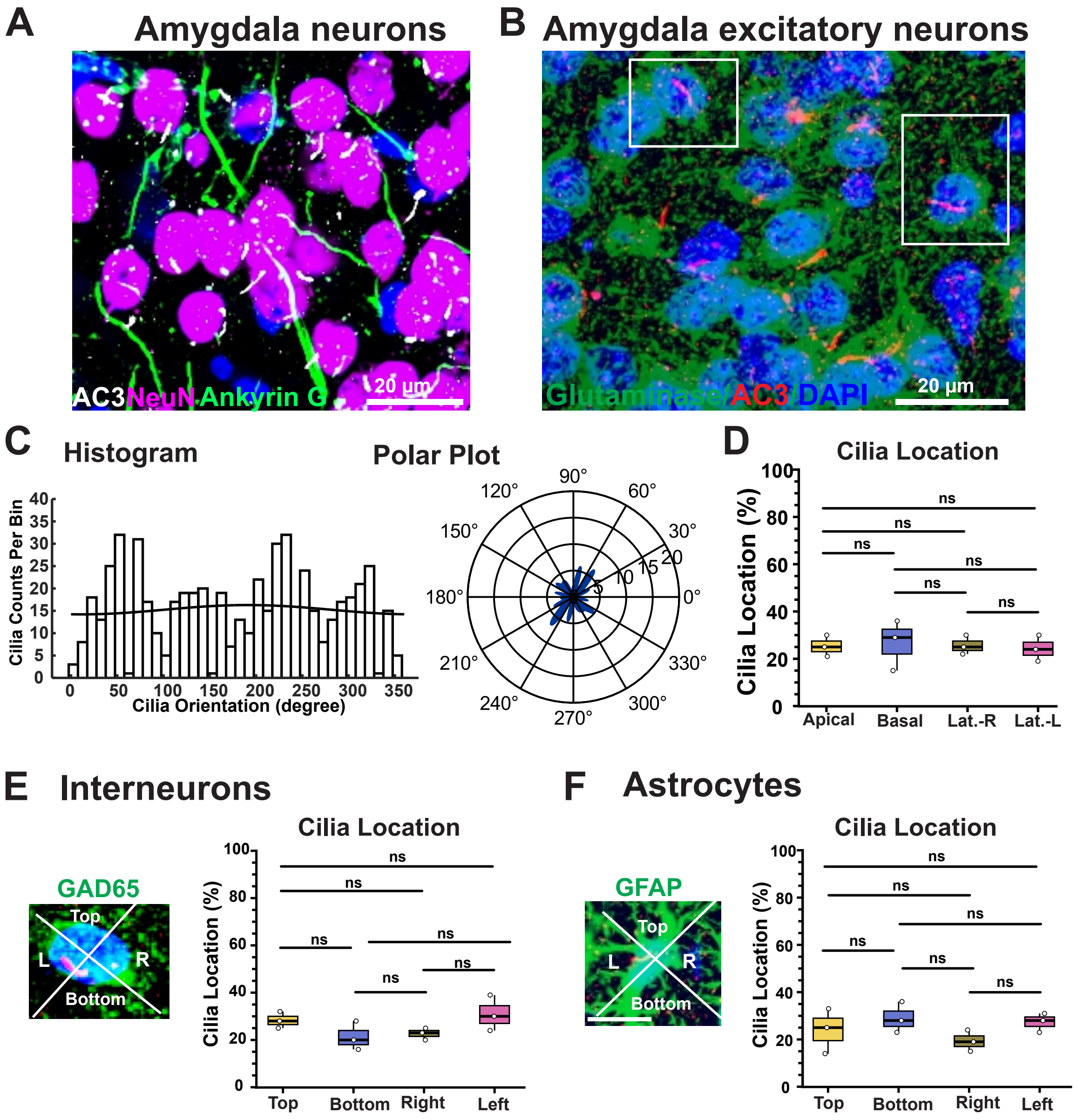

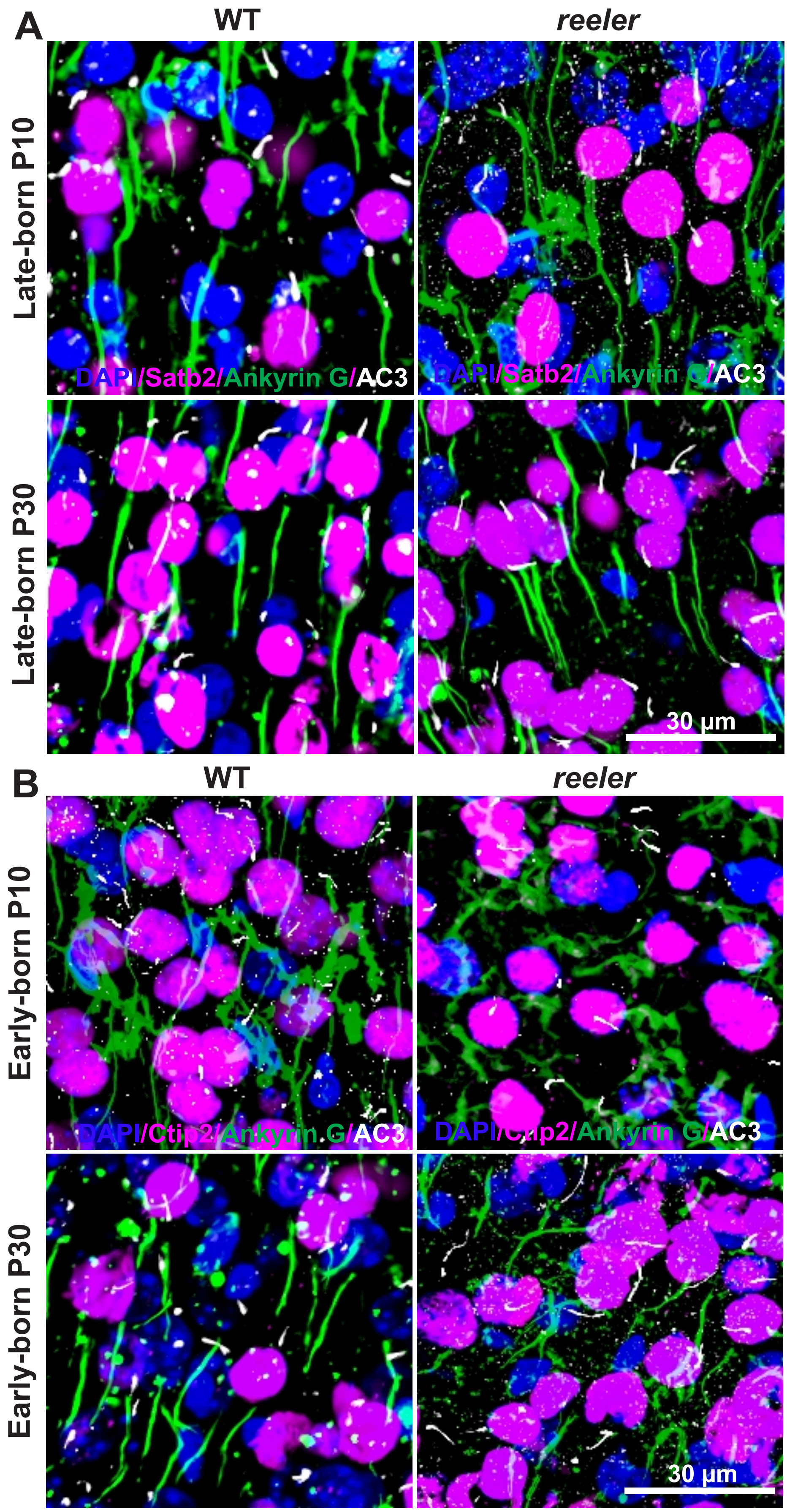

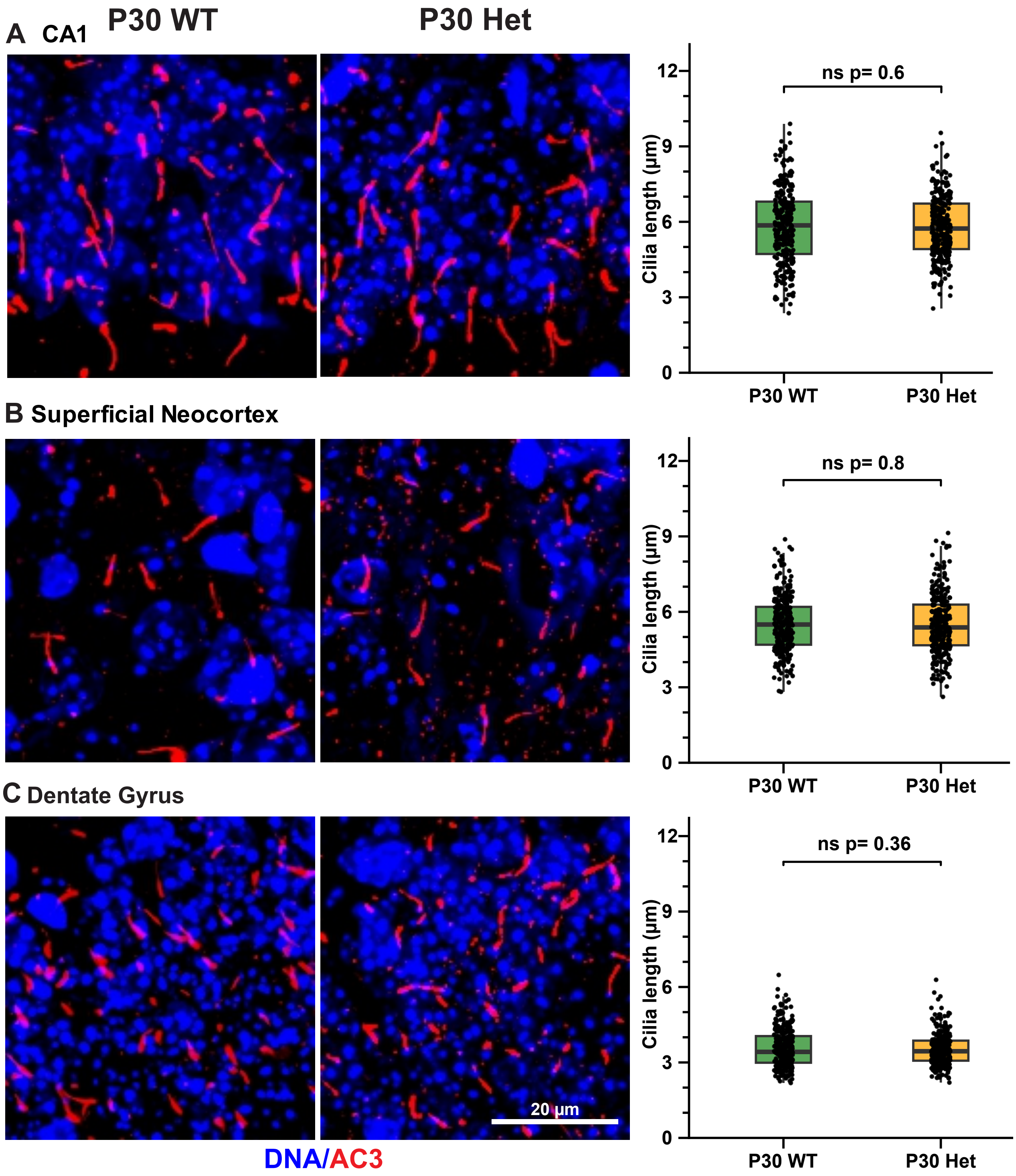

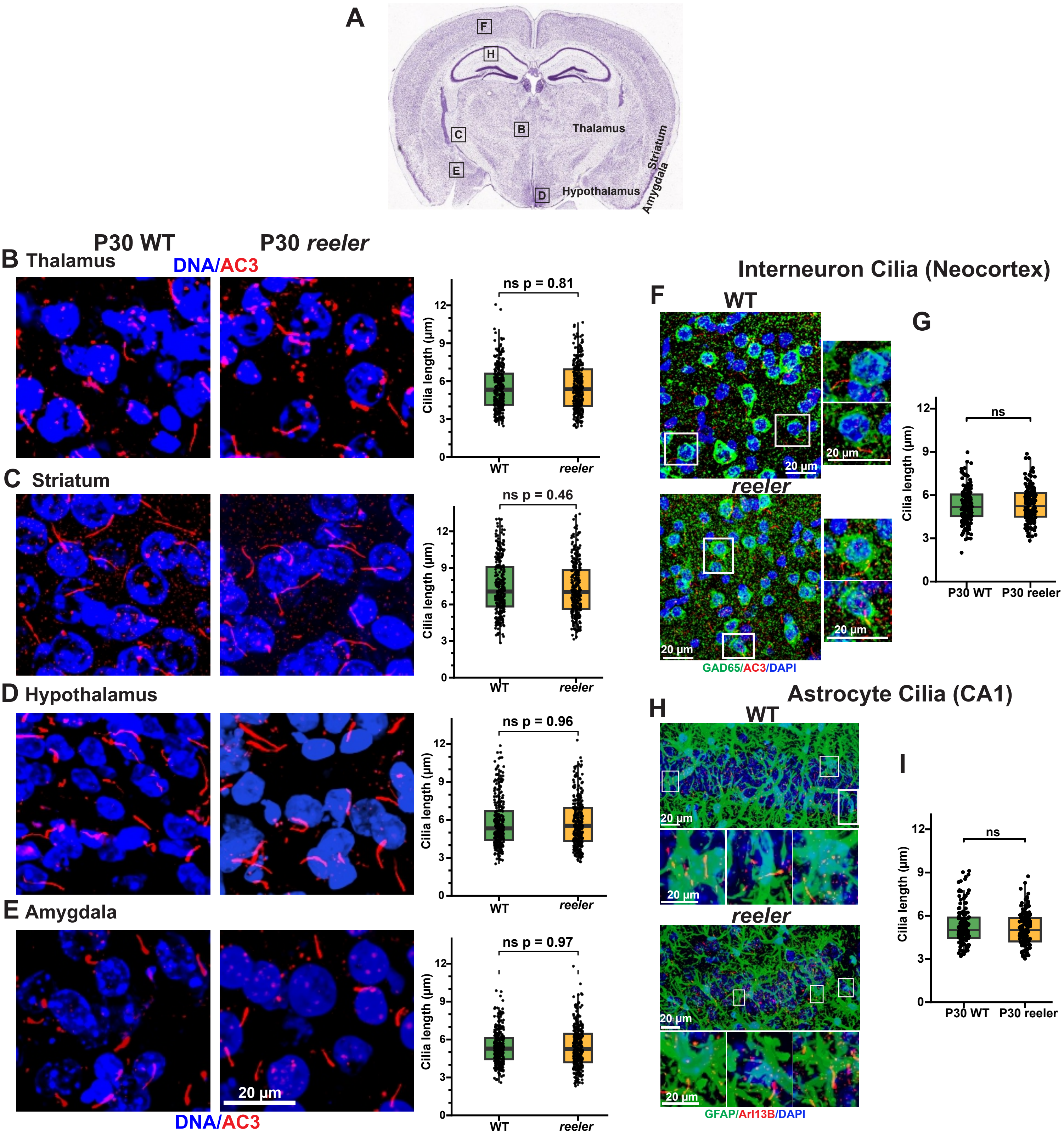
